# Alternative splicing dynamics during human cardiac development *in vivo* and *in vitro*

**DOI:** 10.1101/2025.03.21.642423

**Authors:** Beatriz Gomes-Silva, Marta Furtado, Marta Ribeiro, Sandra Martins, Maria Teresa Carvalho, André Ventura-Gomes, Henrike Maatz, Pragati Nalinkumar Parakkat, Michael Gotthardt, Rosina Savisaar, Maria Carmo-Fonseca

**Author notes:** These authors have contributed equally to the work (Beatriz Gomes-Silva, Marta Furtado). Corresponding author: M. Carmo-Fonseca.

## Abstract

Cardiomyocytes differentiated *in vitro* from human induced pluripotent stem cells (iPSC-CMs) are increasingly used in studies of disease mechanisms, drug development, toxicity testing, and regenerative medicine. Alternative splicing (AS), a crucial mechanism for regulating gene expression during development, plays a pivotal role in cardiac differentiation and maturation. However, the extent to which iPSC-CMs recapitulate native cardiac splicing patterns remains poorly understood. Here, we provide a comprehensive temporal map of AS regulation during human cardiac development. In addition to the major splicing changes occurring perinatally, we identify finely tuned prenatal splicing transitions. iPSC-derived cardiomyocytes globally recapitulate the transcriptome of prenatal cardiomyocytes, yet their splicing profiles remain heterogeneous, with certain events reflecting early embryonic patterns and others resembling those of later-stage heart development. Moreover, we uncover altered splicing events in iPSC-CMs, including mis-splicing of splicing factors. In conclusion, we present a resource of AS dynamics throughout human cardiac development and a catalog of splicing markers to assess cardiomyocyte maturation *in vitro*. Our findings provide critical insights into the limitations of iPSC-CM models and their utility in cardiovascular research.

## INTRODUCTION

Understanding human cardiac development is critical for advancing cardiovascular medicine and uncovering mechanisms underlying congenital and acquired heart diseases. Heart formation relies on the precise temporal control of gene expression programs, involving an evolutionarily conserved network of signaling pathways, transcription factors and epigenetic alterations ^1–4^. These regulatory mechanisms govern the specification of cardiac cell fates and the differentiation and diversification of cardiac cell types throughout development ^5^.

A crucial mechanism that regulates gene expression during development is alternative splicing (AS), a process that expands transcript and protein diversity by generating multiple mRNA isoforms from a single gene ^6,7^. Through AS of nascent transcripts (pre-mRNAs), individual genes produce a variety of mRNA species that may differ in stability, localization, or protein coding capacity ^8–10^. Alternatively spliced protein isoforms may have related, distinct or even opposing functions ^9,10^. Tissue-specific regulation of AS plays a crucial role in defining the identity and function of adult tissues, and its regulation is finely tuned during development ^11–13^. AS plays a pivotal role in cardiac cell differentiation and maturation, with numerous splicing isoforms emerging at different developmental stages ^14–16^. Recently, a cross-species comparison of splicing patterns across pre- and postnatal development of multiple organs revealed that splicing regulation is fundamental for heart development ^17^. Despite these insights, a comprehensive characterization of AS during human cardiac development remains lacking. Furthermore, the physiological roles of developmentally regulated splicing isoforms are still poorly understood, limiting our ability to fully appreciate how splicing transitions contribute to heart formation and function.

Major advances in cardiac developmental biology have resulted from studies in murine models^18^. However, despite their utility, rodent heart architecture and cell function are very different from humans ^19^. The discovery that differentiated adult human somatic cells can be reprogrammed into a pluripotent state ^20^ revolutionized the field, enabling the directed differentiation of pluripotent stem cells into well-defined cardiac lineages *in vitro*. Currently, human induced pluripotent stem cell-derived cardiomyocytes (iPSC-CMs) are widely used as an alternative to animal models for disease modeling, drug discovery, and toxicity screening ^21^. Moreover, iPSC-CM transplantation is being explored as a potential strategy to repopulate damaged myocardial tissue and restore heart function ^22^.

Although iPSC-CMs are generally considered to resemble fetal cardiomyocytes ^21^, a systematic global comparison of splicing programs between iPSC-CMs and human hearts has yet to be performed. Such an analysis is crucial to understanding the extent to which iPSC-CMs recapitulate native cardiac splicing transitions and to identifying potential limitations in their maturation.

In this study, we conducted a comparative analysis of AS patterns in iPSC-CMs and human hearts across developmental stages, from early organogenesis to adulthood. Our findings provide a resource of developmentally regulated AS events, capturing splicing transitions that occur as the heart progresses from embryonic to postnatal life. While splicing profiles in iPSC-CMs globally resemble those of prenatal heart samples, we uncovered a subset of splicing events unique to iPSC-CMs, including mis-splicing of genes involved in RNA processing. Together, these findings provide a comprehensive characterization of AS dynamics during human heart development and establish a catalog of splicing events that can serve as benchmarks for assessing cardiomyocyte maturation *in vitro*.

## RESULTS

### Genome-wide comparison of splicing profiles in iPSC-CMs and developing hearts

To evaluate whether iPSC-CMs recapitulate the splicing patterns of the heart and determine the developmental stage they most closely resemble, we directed the differentiation of three independent iPSC lines (referred to as D, G, and T, **Figure 1A**) into cardiomyocytes (see **Methods**). The majority of differentiated iPSC-CMs exhibited cardiomyocyte-like morphology, including an elongated shape and well-defined sarcomere structures (**Figure 1B, C**).

**Figure 1.**
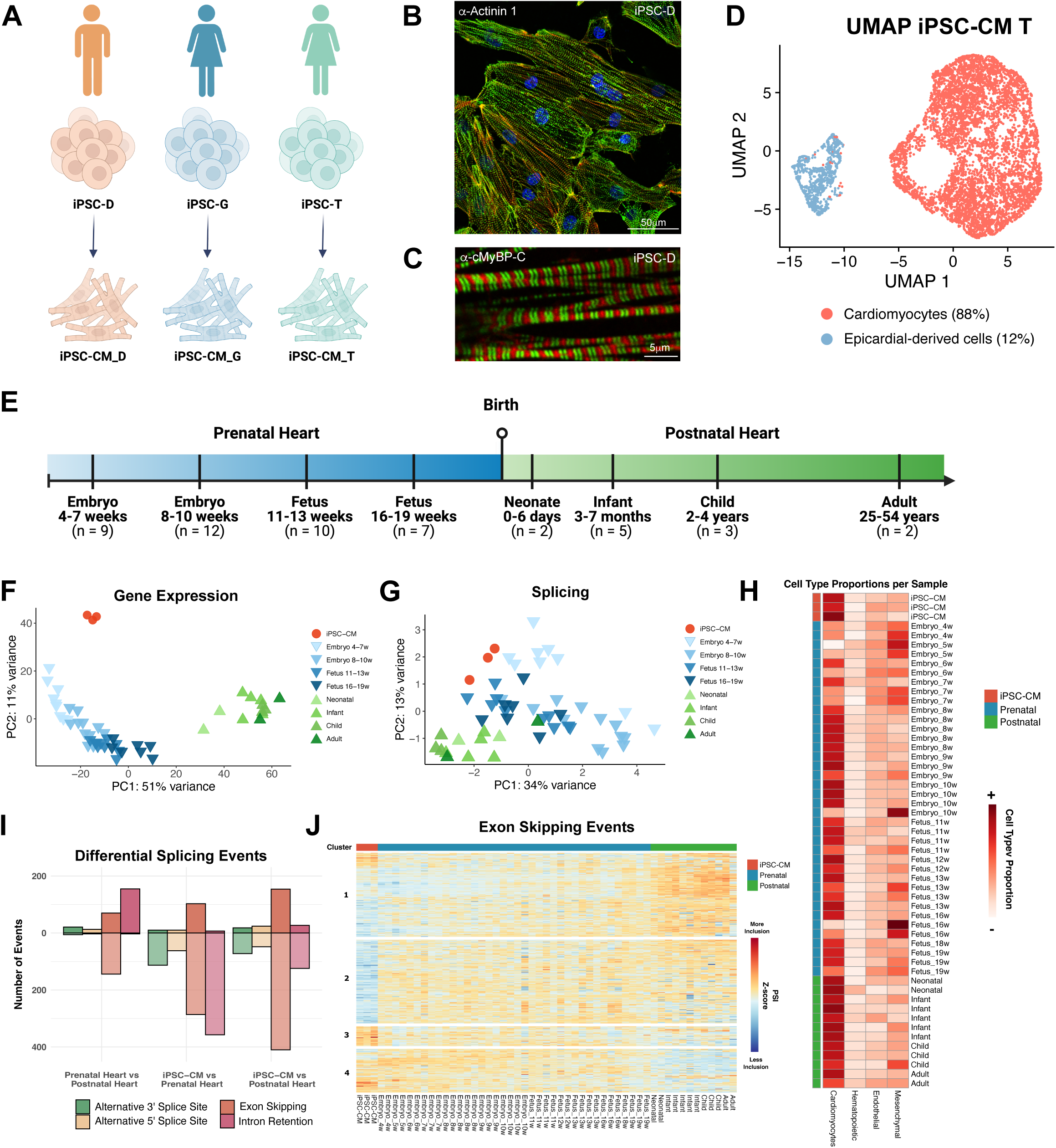
Genome-wide comparison of splicing profiles in iPSC-CMs and developing hearts. **A.** Schematic representation of iPSC differentiation into iPSC-CMs. The iPSC cell lines and their respective sexes are indicated. **B-D.** Immunofluorescence images showing the expression of (**B**) α-Actinin 1 and (**C**) α-MyBP-C, in iPSC-CMs on day 30 of differentiation. Samples were derived from the indicated iPSC lines. Nuclei are stained with DAPI. Scale bar included in each image. **D.** UMAP plot of iPSC-CMs derived from iPSC-T, revealing two major subpopulations of cells corresponding to cardiomyocytes (red), and epicardial-derived cells (blue). **E**. Schematic representation of the developing human heart RNA-seq dataset, with the corresponding timepoints. **F-G.** Principal Component Analysis based on the (**F**) 500 most variable protein-coding genes and (**G**) 500 most variable splicing events, across iPSC-CMs (in red), prenatal hearts (in blue) and postnatal hearts (in green). Lighter shades represent younger samples, and darker shades correspond to older samples. Splicing events in (**G**) include exon skipping, intron retention, and alternative 3’ and 5’ splice sites detected by vast-tools. **H.** Heatmap showing estimated cell type proportions in heart and iPSC-CM samples, determined using CIBERSORTx. **I.** Barplot depicting the total number of differentially spliced events (ΔPSI ≥ ±0.2, adjusted p-value ≤ 0.01) in comparisons between Prenatal vs. Postnatal Heart, iPSC-CM vs. Prenatal Heart, and iPSC-CM vs. Postnatal Heart. Events above 0 (darker color) indicate higher inclusion in the first group of each comparison, while events below 0 (lighter color) indicate lower inclusion in the first group. **J.** Heatmap displaying Z-score normalized Percent Spliced In (PSI) levels for all exon skipping events, comparing iPSC-CMs with prenatal hearts, iPSC-CMs with postnatal hearts, and prenatal hearts with postnatal hearts. Colors reflect inclusion levels relative to the mean of each event (red for higher inclusion, blue for lower inclusion). Event-specific information is provided in Tables S2 and S3.

To assess which cell types were present in monolayer cultures of iPSC-CMs at day 30 of differentiation, we conducted single-cell RNA sequencing (sequencing metrics in **Table S1**). The results revealed that 84–88% of cells expressed gene signatures characteristic of cardiomyocytes (**Figure 1D, Figure S1A**), including the expression of sarcomeric and mitochondrial genes (**Figure S1B, C**). In contrast, ∼12-16% of cells were enriched in *TNNT1*, the predominant troponin isoform during early cardiac development, as opposed to *TNNT2*, which is characteristic of fully mature cardiomyocytes (**Figure S1B**). Label transfer annotation using the Farah *et al*. fetal heart single-cell atlas ^23^ confirmed these cells resembled the gene expression profile of epicardial cells (**Figure S1D**). Accordingly, these cells displayed expression of epicardial markers, including *WT1*, *UPK3B* and *ALDH1A2* (**Figure S1B**) ^24,25^. The epicardial-like cells were enriched in extracellular matrix-related pathways, whereas the cardiomyocyte population showed enrichment in cardiac development pathways (**Figure S1E**). This observation aligns with studies reporting the bifurcation of cardiac progenitors into myocardial and epicardial lineages during both iPSC differentiation ^26^ and mouse cardiac development ^27^.

Next, we conducted bulk RNA-sequencing (RNA-seq) of polyadenylated RNA isolated from the three iPSC-CM lines at day 30 of differentiation (sequencing metrics in **Table S1**). To assess potential variability among iPSC-CMs derived from unrelated individuals, we performed pairwise comparisons of gene expression profiles across the three lines (**Figure S1F**). This analysis revealed a high degree of similarity across the lines, with correlation coefficients comparable or even exceeding those observed between two embryonic heart samples at the same developmental stage (**Figure S1F**), suggesting that the differentiation process yields highly similar iPSC-CMs regardless of donor origin. Additionally, we observed a substantial overlap in expressed genes among all three lines (**Figure S1G**), reinforcing the consistency of the transcriptional landscape across replicates.

To understand how the transcriptomes of iPSC-CMs compare with gene expression profiles during heart development, the three data sets from iPSC-CM cultures were grouped as biological replicates and compared with publicly available bulk RNA-seq datasets from human heart samples, spanning from early organogenesis to adulthood ^28^. This dataset comprises 38 prenatal samples collected between 4 and 19 weeks post-conception, and 12 postnatal samples collected from newborns, infants, young children, and adults (**Figure 1E**). To explore the primary sources of variation in gene expression profiles of iPSC-CMs and human hearts at distinct developmental stages, we performed principal component analysis (PCA). This analysis revealed that the first principal component (PC1), accounting for 51% of the variance, effectively distinguishes prenatal heart samples from postnatal samples, capturing progressive transcriptional changes throughout development (**Figure 1F**). Along PC1, the iPSC-CMs clustered most closely with 8– 10 week prenatal heart samples (**Figure 1F**). Hierarchical clustering revealed that the iPSC-CMs cluster closer to the prenatal heart than to the postnatal heart samples, and bootstrapping confirmed the reliability of the clustering, attributing 100% confidence (approximately unbiased p-value * 100) to the node that clusters the iPSC-CMs with the prenatal hearts (**Figure S1H**).

To further assess the developmental state of iPSC-CMs, we leveraged our RNA-seq data to examine the expression of genes associated with key metabolic pathways involved in cardiomyocyte maturation. Metabolic maturation of is characterized by shift from anaerobic glycolysis to mitochondrial oxidative phosphorylation (OXPHOS), accompanied by increased fatty acid β-oxidation (FAO) and mitochondrial biogenesis ^29^. Using single-sample Gene Set Enrichment Analysis (ssGSEA), we confirmed that prenatal heart samples showed higher glycolytic activity and lower OXPHOS, while postnatal hearts showed the reverse pattern, consistent with metabolic remodeling (**Figure S1I**). iPSC-CMs retained a glycolytic, prenatal-like metabolic profile (**Figure S1I**). To further refine this analysis, we curated cardiomyocyte-relevant gene panels for Glycolysis, the TCA cycle, OXPHOS, and FAO, and visualized their expression in a combined heatmap (**Figure S1J**). While iPSC-CMs express key components of the TCA cycle and OXPHOS machinery, similar to late-prenatal hearts, the expression of FAO-related genes remained markedly low, even compared to early-prenatal hearts (**Figure S1J**). These findings align with prior studies ^30,31^ and suggest that although iPSC-CMs possess functional mitochondria capable of oxidative phosphorylation, they fail to activate the FAO program, a hallmark of metabolically mature cardiomyocytes.

To identify and quantify alternative splicing (AS), we used rMATS, MAJIQ, and vast-tools, focusing on the most common patterns of AS events (i.e., exon skipping, alternative 3’ and 5’ splice sites, and intron retention). Pairwise comparisons of the inclusion levels of all AS events detected in the three iPSC-CM cultures revealed Spearman correlation coefficients of ∼0.93, higher than that observed between embryonic hearts at the same developmental age (**Figure S1K**). The three iPSC-CM datasets were grouped and compared with heart datasets. While PCA of splicing data was noisier than that of gene expression, considering that developmental time is captured by the combination of the two principal components, iPSC-CMs consistently clustered closer to the prenatal samples, across all three tools (**Figure 1G**, **Figure S1L, M)**. Given the distinct cell type composition of iPSC-CMs and heart tissues, differences in splicing patterns could be influenced by sample heterogeneity. To explore this possibility, we applied CIBERSORTx to estimate the relative abundance of major cardiac cell types in our bulk RNA-seq datasets using a single-cell reference. This analysis revealed that iPSC-CMs were highly enriched in cardiomyocytes, consistent with their purification by fluorescence-activated cell sorting with VCAM1 antibodies at day 13 of differentiation, whereas prenatal heart samples exhibited greater cellular diversity and a comparatively lower proportion of cardiomyocytes (**Figure 1H**). While CIBERSORTx does not provide definitive cell-type quantification, it offers a useful approximation to assess whether compositional differences might influence transcriptome-wide splicing patterns. As the cardiomyocyte content of iPSC-CMs was more similar to postnatal than to prenatal samples (**Figure 1H**), it is thus unlikely that their transcriptional clustering with prenatal hearts is solely driven by cell type composition.

Pairwise differential splicing analysis, focusing on identifying events with a deltaPSI (difference in Percent Spliced In) higher than 20%, revealed 411 differentially spliced events between prenatal and postnatal heart samples (**Figure 1I, Table S2**). When comparing iPSC-CMs and heart samples, we found 948 splicing events differing between *in vitro* cultures and prenatal samples, and 877 events dissimilar between *in vitro* differentiated cardiomyocytes and postnatal heart samples (**Figure 1I, Table S3**). Alternative ‘‘cassette’’ exon and intron retention were the most frequent types of alternative splicing identified (**Figure 1I**).

Hierarchical clustering of differentially spliced events revealed distinct patterns of exon usage and intron retention across developmental stages. Exons predominantly skipped in prenatal hearts were grouped in cluster 1, while those primarily skipped in postnatal hearts were grouped in cluster 4 (**Figure 1J**). Notably, within cluster 1, the AS events in iPSC-CMs segregated into two subgroups: a smaller upper subgroup with splicing patterns more closely resembling postnatal hearts, and a larger lower subgroup aligning more closely with prenatal profiles, highlighting the heterogeneous nature of AS in iPSC-CMs. The analysis also identified exon skipping events that uniquely distinguish iPSC-CMs from both prenatal and postnatal hearts (**Figure 1J** – clusters 2 and 3), as well as intron retention events specific to iPSC-CMs (**Figure S1N** - cluster 2). Finally, a skew toward increased intron retention was observed in prenatal hearts (**Figure S1N** – cluster 3).

Taken together, our findings indicate that iPSC-CMs globally resemble prenatal cardiac cells at both the transcriptional and splicing levels, although some variability is observed across individual splicing events.

### Pre-to postnatal splicing transitions in the heart

Among the 411 events differentially spliced between prenatal and postnatal heart samples (**Table S2**), most had not been previously reported as developmentally regulated during human cardiac maturation *in vivo* and were predicted to have functional consequences at the protein level (**Table S4**). To infer the cell-type specificity of the genes affected by these splicing events, we analyzed their expression using published single-cell RNA-seq data from human hearts ^23^. This analysis showed that many of the newly identified events occur in genes that are either predominantly expressed in non-cardiomyocyte populations or are broadly expressed across multiple cardiac cell types. Thus, to avoid potential misinterpretation due to differences in cellular composition, we excluded these events from comparisons with iPSC-CM samples.

To investigate the potential functional significance of the newly identified events, we used *VastDB* and *NEASE* to predict their effects on protein function and interactions. A striking enrichment of splicing events predicted to impact proteins associated with the extracellular matrix (ECM) was observed (**Figure 2A**). Among affected ECM genes, we observed alternative splicing of fibronectin (*FN1*) exons 25 (**Figure 2B**) and 33 (**Figure S2A**), corresponding to the EDB and EDA isoforms ^32^. We also observed AS changes in Kindlin-2 (*FERMT2*), a regulator of integrin-mediated cell adhesion and ECM remodeling ^33^. In the postnatal heart, we found increased inclusion of exon 11 (**Figure 2C**), which encodes a domain involved in various protein interactions. Relative to pre-natal hearts, we also found increased inclusion of Fibulin-2 (*FBLN2*) exon 9 (**Figure S2B**), which encodes a calcium-binding EGF domain ^34^.

**Figure 2.**
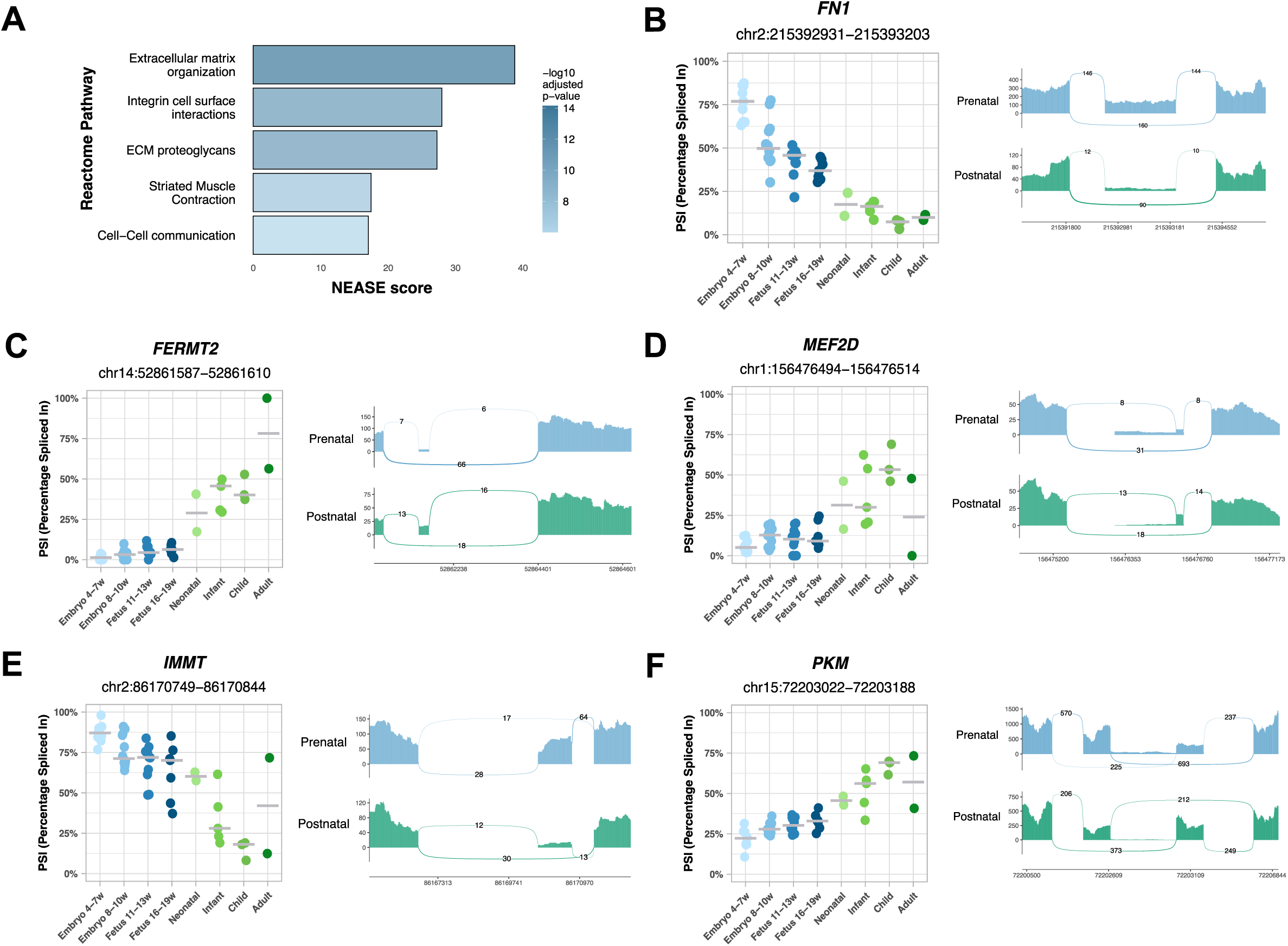
Pre-to postnatal splicing transitions in the heart. **A.** Top 5 enriched Reactome pathways identified by NEASE for differentially spliced exons identified between prenatal and postnatal hearts. **B-F.** Inclusion levels (PSI) of exons differentially included between prenatal and postnatal hearts, specifically (**B**) *FN1* exon 25 (*FN1*-203), (**C**) *FERMT2* exon 11 (*FERMT2*-215), (**D**) *MEF2D* exon 8 (*MEF2D*-201), (**E**) *IMMT* exon 6 (*IMMT*-204), and (**F**) *PKM* exon 9 (*PKM*-219). PSI levels are shown for prenatal hearts and postnatal hearts. Each event is accompanied by the corresponding sashimi plot.

We further detected splicing changes in the transcription factor *MEF2D*, with increased inclusion of exon 8, which encodes a domain essential for transcriptional activation ^35^, occurring after birth (**Figure 2D**). Mutually exclusive alternative splicing produces two major *TCF3* isoforms which differ only in their DNA binding domain ^36^ During heart development, the isoform E47-specific exon was more included after birth (**Figure S2C**). Increased inclusion of *MLF1* exon 3, which disrupts a domain required for cell cycle control ^37^ was further detected in postnatal hearts (**Figure S2D**).

Additionally, we identified AS events in metabolism-related genes. We observed a progressive decrease in inclusion of *IMMT* exon 6 during development, particularly after birth (**Figure 2E**). *IMMT* encodes the mitochondrial protein mitofilin ^38^, and its exon 6 encodes a domain required for the function of mitochondrial cristae ^39^. We also detected an AS switch in *PKM* (**Figure 2F**). This gene encodes pyruvate kinase M, an enzyme that catalyzes the final step in glycolysis. Exons 9 and 10 of *PKM* are mutually exclusive, giving rise to two isoforms with distinct metabolic roles: exon 9 inclusion produces PK-M1, an isoform predominantly expressed in terminally differentiated tissues with high oxidative demands, while exon 10 inclusion generates PK-M2, an isoform commonly found in proliferative fetal tissues and cancer ^40^. Our finding that *PKM* exon 9 inclusion is higher in postnatal hearts compared to prenatal tissues (**Figure 2F**) suggests that this splicing transition contributes to the metabolic shift from glycolysis to oxidative phosphorylation during cardiac maturation after birth.

In summary, we identified numerous novel cardiac AS events that are regulated during the transition from pre-to postnatal life. These splicing alterations are predicted to affect critical cardiac functions, including extracellular matrix remodeling, transcriptional control, and metabolic adaptation.

### Splicing regulation in the developing prenatal heart

While the transition to postnatal life is a critical point for splicing regulation, splicing patterns during the prenatal period are not uniform across developmental stages. A comparison of early embryonic (4-5 weeks post-conception) and later fetal (18-19 weeks post-conception) heart samples revealed 563 alternative exon skipping events and 73 intron retention events (**Table S5**).

Embryonic hearts showed higher levels of exon skipping (**Figure 3A)** and intron retention (**Figure S3A**) compared to fetal hearts. Genes undergoing differential splicing during the embryonic to fetal transition were most strongly enriched in pathways associated with membrane trafficking, cell-cell communication and VEGF signaling (**Figure 3B, Table S6**).

**Figure 3.**
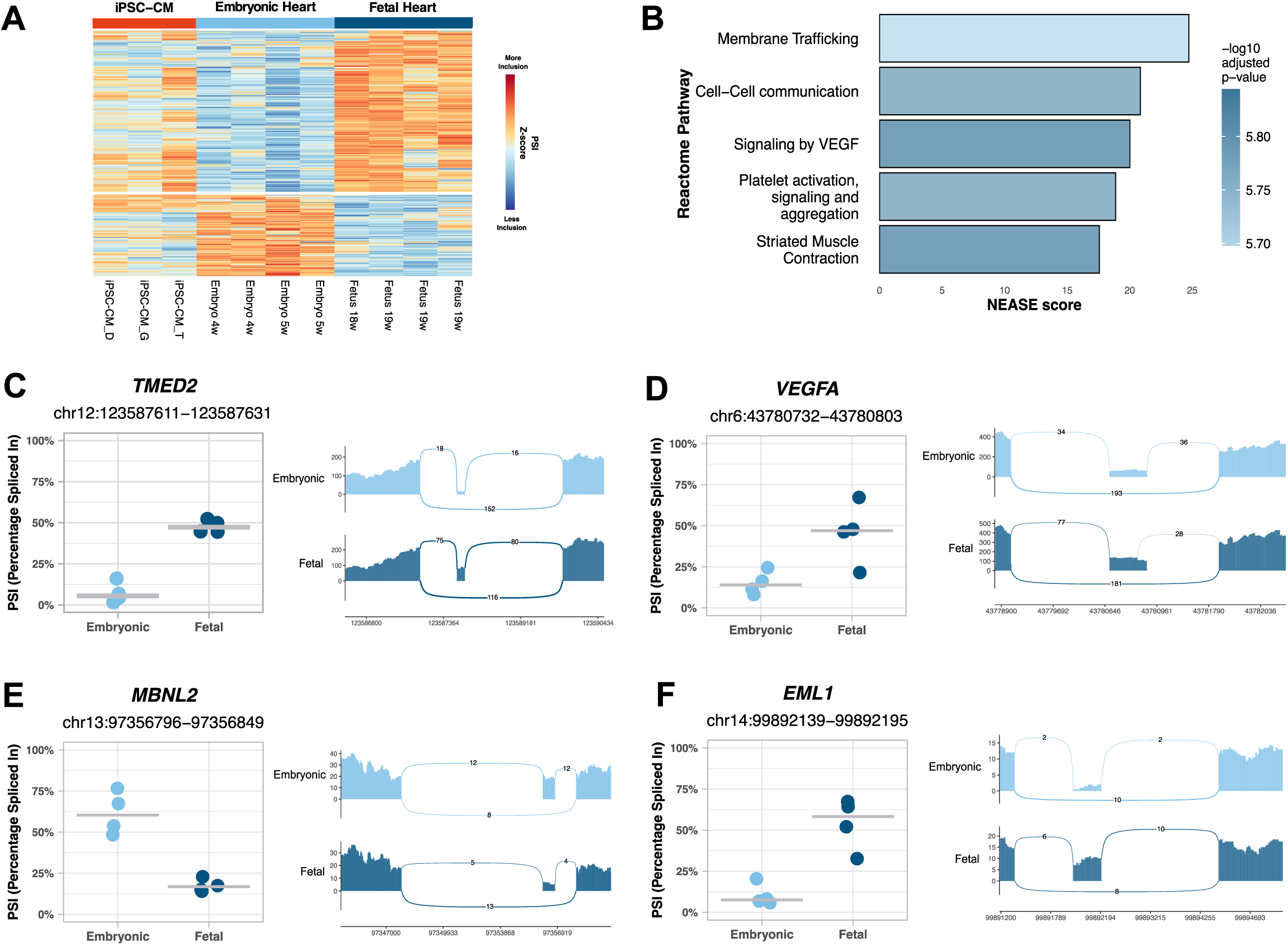
Splicing regulation in the developing prenatal heart. **A.** Heatmap displaying Z-score normalized Percent Spliced In (PSI) levels of all exon skipping events identified when comparing 4-5 weeks embryonic with 18-19 weeks fetal hearts. Colors represent inclusion levels relative to the mean for each event (red indicates higher inclusion, blue indicates lower inclusion). Event-specific information is provided in Table S5. **B.** Top 5 enriched Reactome pathways identified by NEASE for differentially spliced exons identified between 4-5 weeks embryonic and 18-19 weeks fetal hearts. **C-F**. Inclusion levels (PSI) of exons differentially spliced between 4-5 weeks embryonic and 18-19 weeks fetal hearts, specifically (**C**) *TMED2* exon 3 (*TMED2*-202), (**D**) *VEGFA* exon 6 (*VEGFA*-226), (**E**) *MBNL2* exon 6 (*MBNL2*-207), and (**F**) *EML1* exon 2 (*EML1*-222). PSI levels are shown for embryonic hearts and fetal hearts. Each event is accompanied by the corresponding sashimi plot.

For instance, *TMED2*, a gene involved in cargo selection and vesicle formation and essential for development ^41^, begins to include its heart-specific micro-exon 3 in the fetal stage ^17^ (**Figure 3C)**.

*VEGFA* exon 6, located in the heparin-binding domain, was more included in fetal compared to embryonic hearts (**Figure 3D**).

*MBNL2* exon 6, which encodes a nuclear localization signal, showed higher inclusion in embryonic hearts compared to fetal hearts (**Figure 3E**).

*EML1* exon 2, which encodes a motif thought to mediate interactions with microtubules and membranes, was more included in fetal hearts compared to embryonic hearts (**Figure 3F**).

Additional genes with significant regulation of prenatal splicing included *MACF1*, *LAMA2*, *ANK3*, *ABI1*, *DCAF6*, *SLC25A3* and *ATP2B4* (**Figure S3B-H, Table S6**).

In conclusion, our findings demonstrate that cardiac splicing is dynamically regulated throughout prenatal development.

### Heterogeneous recapitulation of developmental splicing regulation in iPSC-CMs

Although PCA analysis segregated embryonic and fetal hearts along PC2, with iPSC-CMs clustering in between (**Figure 4A**), direct comparisons are confounded by the cellular heterogeneity of prenatal heart tissue, which contains a substantial proportion of non-cardiomyocytes. To mitigate this limitation, we restricted the analysis to genes predominantly expressed in cardiomyocytes. Based on single-cell transcriptomic data from the fetal heart atlas ^23^, we identified 491 cardiomyocyte-enriched genes (log2FC > 0.5 compared to all other cell types), of which 86 met stricter criteria for specificity (expression in >50% of cardiomyocytes and <10% of non-cardiomyocytes). Within these cardiomyocyte-enriched genes (**Table S7; Figure 4B, S4A**), we detected 79 AS events regulated during prenatal development. Comparison of these events revealed broad variability between iPSC-CMs and prenatal hearts. In some cases, iPSC-CM splicing patterns closely mirrored those of embryonic hearts (**Figure 4C-F, S4B-D**), whereas other events were more similar to fetal hearts (**Figure 4G, S4E-G**) or displayed intermediate profiles (**Figure 4H-J**). Strikingly, even within the same gene, different AS events could align with distinct developmental stages (**Figure S4H-O**). For a subset of AS events, the pattern in iPSC-CMs diverged further from that of embryonic and fetal hearts (**Figure S4P-Q**), suggesting a more mature splicing profile *in vitro*. Together, these findings indicate that iPSC-CMs display a heterogeneous maturation state with respect to developmental splicing regulation in the prenatal heart.

**Figure 4.**
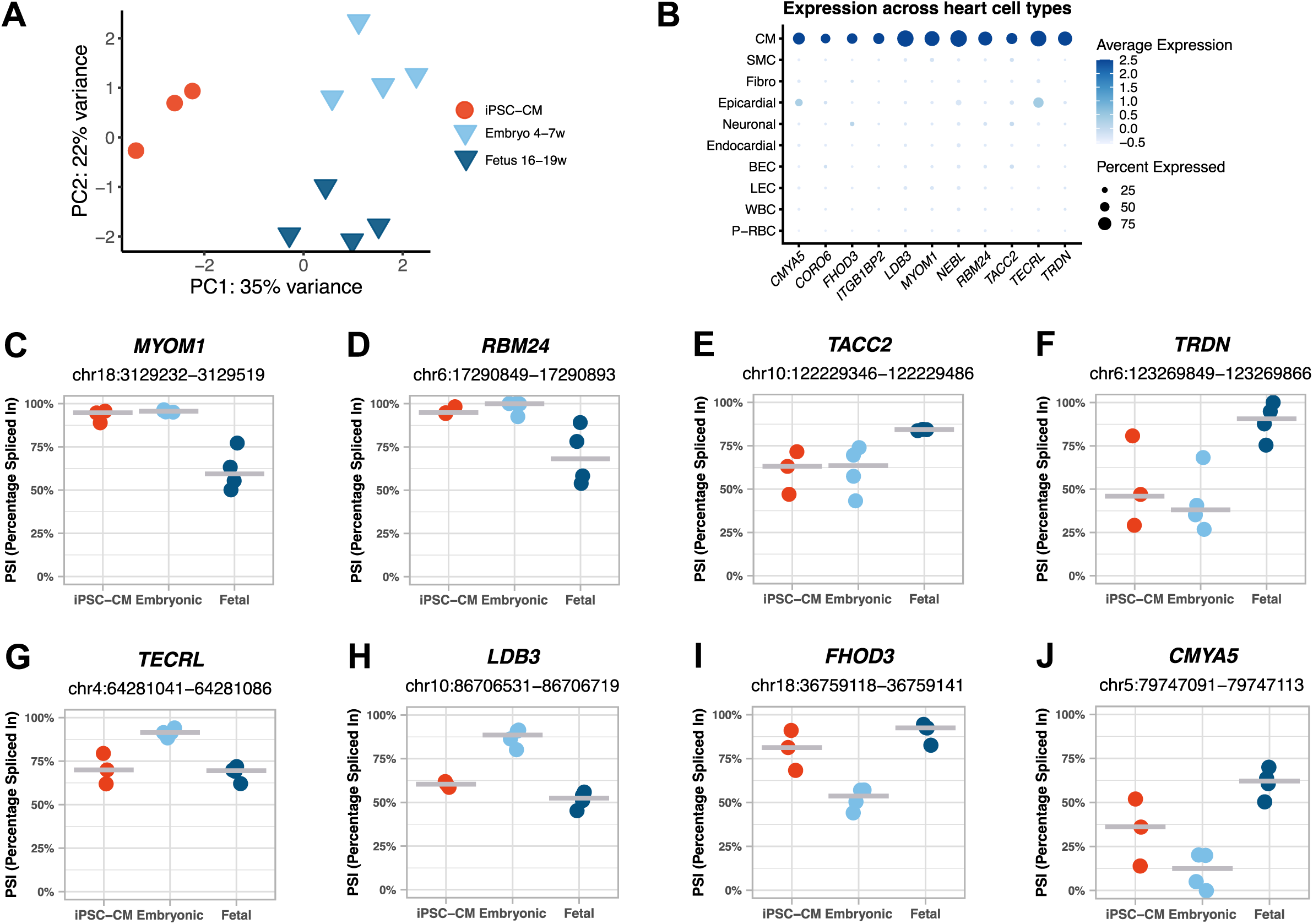
Heterogeneous mirroring of prenatal heart splicing regulation in iPSC-CMs. **A.** PCA of the 500 most variable splicing events in iPSC-CMs (red), 4-5 weeks embryonic hearts (in light blue) and 18-19 weeks fetal hearts (in dark blue), as identified by MAJIQ. **B.** Dot plot depicting the expression of cardiomyocyte-specific genes (log2FC > 0.5 compared with all other cell types, expressed in >50% of cardiomyocytes and <10% of other cell types) across the fetal heart single-cell atlas. Dot size represents the percentage of cells expressing each gene, and dot color indicates the scaled expression levels averaged per cell population. **C-J**. Inclusion levels (PSI) of exons differentially spliced between 4-5 weeks embryonic and 18-19 weeks fetal hearts, specifically (**C**) *MYOM1* exon 18 (*MYOM1*-202), (**D**) *RBM24* exon 4 (*RBM24*-207), (**E**) *TACC2* exon 15 (*TACC2*-210), (**F**) *TRDN* exon 31 (*TRDN*-201), (**G**) *TECRL* exon 11 (*TECRL*-201), (**H**) *LDB3* 8 (*LDB3*-202), (**I**) *FHOD3* exon 26 (*FHOD3*-206), and (**J**) *CMYA5* exon 5 (*CMYA5*-201). PSI levels are shown for iPSC-CMs, embryonic hearts, and fetal hearts.

### Maturation-driven splicing changes in iPSC-CMs recapitulate developmental transitions in the human heart

The observation that iPSC-CMs mimic prenatal heart splicing patterns but vary in their resemblance to specific developmental stages suggests heterogeneity in splicing trajectories during *in vitro* differentiation. To investigate this, we selected five splicing events for analysis by qRT-PCR at different stages of iPSC-CM maturation. We designed two experimental setups to assess how differentiation time and mechanical environment influence splicing regulation in iPSC-CMs. In one set of experiments, iPSC-CMs differentiated for 30 days were maintained in culture for an additional 10 days (**Figure 5A**). In another set of experiments, iPSC-CMs differentiated for 30 days were cultured for an additional 10 days on a micropatterned surface (**Figure 5A**). We analyzed cardiomyocytes derived from three independent iPSC lines, performing three separate differentiation experiments for each line. As a hallmark of maturation, we assessed sarcomere length (**Figure 5B**). We observed a significant increase in sarcomere length between day 30 and day 40 (**Figure 5B**). Except for iPSC-CM C, culture on a micropatterned surface further increased the sarcomere length (**Figure 5B**).

**Figure 5.**
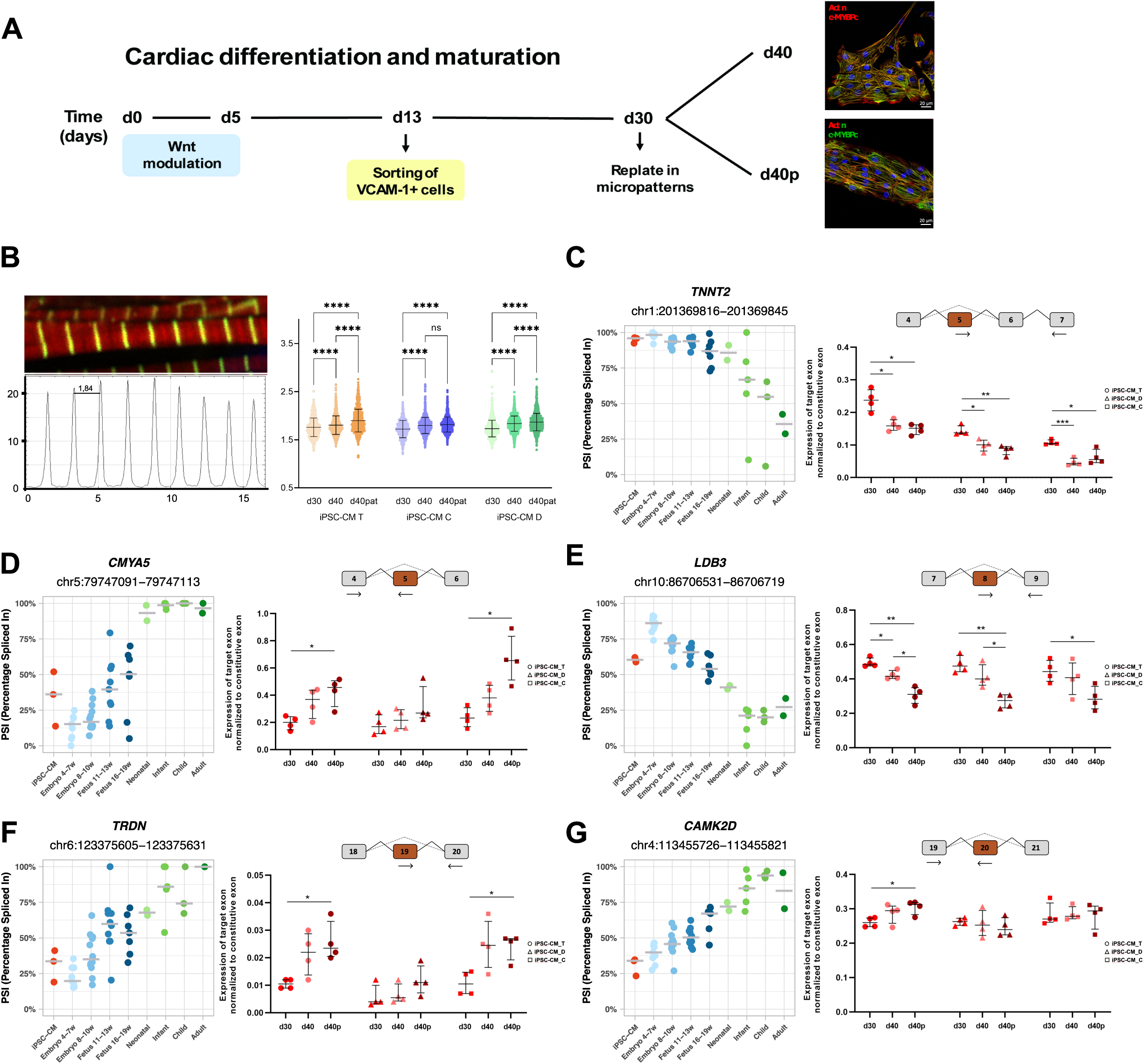
Maturation-driven splicing changes in iPSC-CMs recapitulate developmental transitions in the human heart. **A**. Diagram illustrating the differentiation protocol used to generate iPSC-CMs and the experimental conditions for assessing alternative splicing during *in vitro* maturation. Conditions include iPSC-CMs at 30 days of differentiation, 40 days of differentiation, and 40 days of differentiation with micropatterning. **B.** Immunofluorescence image showing expression of α-Actinin 1 (green) and Actin (red) in sarcomeres of iPSC-CMs at day 30 of differentiation. Plot profile of α-Actinin 1 fluorescence intensity across sarcomeres. Sarcomere length measurements of iPSC-CMs derived from three cell lines (T, D, C) across three experimental conditions. **C-G.** Inclusion levels (PSI) of exons differentially spliced between prenatal and postnatal hearts, specifically (**C**) *TNNT2* exon 5 (*TNNT2*-224), (**D**) *CMYA5* exon 5 (*CMYA5*-201), (**E**) *LDB3* exon 8 (*LDB3*-202), (**F**) *TRDN* exon 19 (*TRDN*-201) and (**G**) *CAMK2D* exon 18 (*CAMK2D*-202). PSI levels are shown for iPSC-CMs, prenatal hearts, and postnatal hearts across various developmental time points. Each splicing event is accompanied by *in vitro* PCR quantification performed in three distinct cell lines (T, D and C) under three experimental conditions: 30 days of differentiation (in red), 40 days of differentiation (in pink), and 40 days of differentiation with micropatterning (in dark red).

For *TNNT2* exon 5, which increases calcium sensitivity and is predominantly included in prenatal hearts ^42^, we observed a reduction in inclusion levels at day 40 compared to day 30. Culturing iPSC-CMs on a micropatterned surface did not further influence exon 5 exclusion (**Figure 5C**).

In *CMYA5*, which encodes a protein required for cardiac dyad architecture ^43^, we identified novel exon skipping and intron retention events in prenatal hearts (**Figure 5D, S5A**) that are predicted to disrupt the reading frame and trigger nonsense-mediated decay. Consistent with this prediction, the levels of *CMYA5* mRNA were significantly reduced in prenatal hearts (**Figure S5B**). In hearts, inclusion of exon 5 increased progressively during prenatal development (**Figure 5D**). In iPSC-CMs, we observed a similar trend, with exon 5 inclusion increasing between day 30 and day 40 of differentiation. This effect was particularly pronounced in cells cultured on a micropatterned surface (**Figure 5D**).

We identified several exons in the sarcomeric Z-line protein gene *LDB3* whose inclusion is developmentally regulated in the heart (**Figure 5E**, **Figure S5C-E**) ^44^. Inclusion of exon 8 decreased progressively during prenatal development (**Figure 5E**). A significantly reduced inclusion of this exon was also observed in iPSC-CMs between day 30 and day 40, and this effect was more pronounced in iPSC-CMs cultured on micropatterned surfaces (**Figure 5E**).

For *TRDN*, a key regulator of calcium release ^45^, we identified a progressive increase in exon 19 inclusion during prenatal heart development (**Figure 5F**). A similar trend of increased inclusion was observed in iPSC-CMs between day 30 and day 40 (**Figure 5F**).

For *CAMK2D*, a gene involved in calcium handling and transcriptional regulation ^46^, exon 18 inclusion increased progressively during heart development (**Figure 5G**). Surprisingly, we detected little to no modulation of exon 18 inclusion in iPSC-CMs between day 30 and day 40, regardless of whether the cells were cultured on micropatterned surfaces (**Figure 5G**). These findings suggest that the regulation of exon 18 inclusion in *CAMK2D* during development is not replicated in iPSC-CMs.

In conclusion, our results demonstrate that, despite inherent variability across iPSC lines and differentiation experiments, both temporal progression in culture and mechanical cues from micropatterned surfaces significantly influence splicing regulation. Except for *CAMK2D*, we observed a consistent trend of recapitulating the developmental splicing transitions characteristic of *in vivo* heart development.

To evaluate whether developmentally regulated alternative transcripts are translated into distinct protein isoforms, we performed deep proteomic profiling of iPSC-CMs. Two of the iPSC lines used for proteomic analysis were the same as those profiled by RNA-seq (T and D); however, iPSC line G was replaced by line C due to sample availability. All iPSCs were differentiated to cardiomyocytes using the same protocol. In parallel, we analyzed previously published high-resolution proteomic datasets from healthy adult human hearts ^47^. Our strategy for detecting translated isoforms focused on AS events that are developmentally regulated and occur in genes predominantly expressed in cardiomyocytes (**Table S8**). As an illustrative example, AS of exon 18 in the *MYOM1* gene results in isoform-specific peptides: exon inclusion gives rise to peptides spanning exons 17–18 and 18–19, while exon skipping produces a junctional peptide spanning exons 17–19. In iPSC-CMs, we detected 4 MS/MS spectra supporting translation of isoforms resulting from exon inclusion. In adult heart samples, peptides corresponding to both isoforms were detected, with a predominance of exon-skipping peptides (**Table S8**). This is consistent with the RNA-seq observation that exon 18 is largely excluded from *MYOM1* transcripts in the adult heart.

Additional AS events identified in our transcriptomic analysis and supported by isoform-specific peptide evidence are summarized in **Table S8**.

### A subset of splicing events distinguishes iPSC-CMs from hearts

In addition to *CAMK2D* exon 18, whose inclusion is developmentally regulated in the heart but remains largely unchanged during iPSC-CM maturation (**Figure 5G**), we identified a subset of splicing events that display distinct patterns in iPSC-CMs compared to human hearts, irrespective of developmental stage (**Figure 1J** - clusters 2 and 3**, Figure S1N** - cluster 2). Our statistical analysis revealed 333 exons with splicing patterns unique to iPSC-CMs (**Table S9**). These events were enriched in genes associated with striated muscle contraction and RNA metabolism, including mRNA splicing (**Figure 6A, Table S10**).

**Figure 6.**
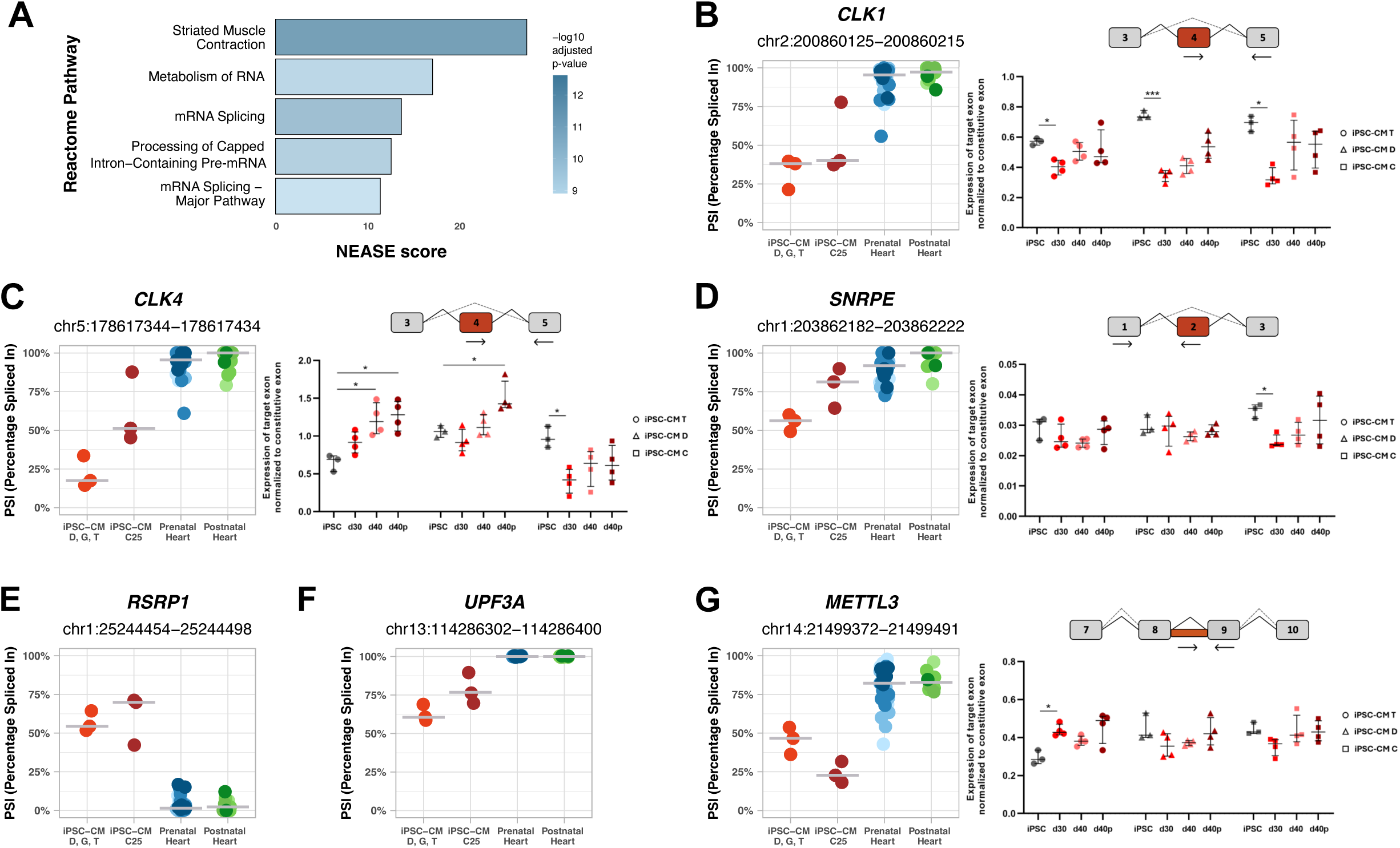
Splicing events with divergent patterns in iPSC-CMs and hearts. **A.** Top 5 enriched Reactome pathways identified by NEASE for differentially spliced exons identified between iPSC-CMs and both prenatal and postnatal hearts. **B-G**. Inclusion levels (PSI) of exons/introns differentially spliced between iPSC-CMs and both prenatal and postnatal hearts, specifically (**B**) *CLK1* exon 4 (*CLK1*-201), (**C**) *CLK4* exon 4 (*CLK4*-201), (**D**) *SNRPE* exon 2 (*SNRPE*-202), (**E**) *RSRP1* exon 4 (*RSRP1*-*204)*, (**F**) *UPF3A* exon 4 (*UPF3A*-202), and (**G**) *METTL3* intron 8 (*METTL3*-201). PSI levels are shown for iPSC-CMs purified by fluorescence-activated cell sorting with VCAM1 antibodies (iPSC-CMs D, G, T), iPSC-CMs purified by metabolic selection with lactate (iPSC-CM C25), prenatal hearts, and postnatal hearts. (**B**), (**C**), (**D**) and (**G**) are accompanied by *in vitro* PCR quantification performed in three distinct cell lines (T, D and C) under four experimental conditions: iPSC not differentiated (in grey), 30 days of differentiation (in red), 40 days of differentiation (in pink), and 40 days of differentiation with micropatterning (in dark red).

Many splicing events that differ between iPSC-CMs and heart tissue occur in genes not exclusively expressed in cardiomyocytes, raising the possibility that differences in cell-type composition could confound interpretation. In heart samples, however, these events were typically constitutively spliced, with PSI values approaching 0 or 1, indicating near-complete exon exclusion or inclusion across developmental stages (**Figure 6B-G**). In contrast, iPSC-CMs displayed intermediate PSI values for the same events, reflecting greater splicing variability (**Figure 6B-G**). These results suggest that the observed splicing differences in iPSC-CMs are unlikely to result from cellular heterogeneity, but rather represent intrinsic features of iPSC-CM biology or the *in vitro* differentiation environment, which diverges from the splicing regulation observed in native heart tissue.

To assess whether iPSC-CM–specific splicing events could result from variability introduced by the differentiation protocol, we expanded our analysis to include RNA-seq datasets from iPSC-derived cultures enriched for cardiomyocytes by metabolic selection with lactate (iPSC-CM C25), in addition to cultures purified by fluorescence-activated cell sorting with VCAM1 antibodies (iPSC-CM C, D, G, T) (see **Methods**). Overall, iPSC-CMs generated with either protocol displayed similar splicing patterns across developmentally regulated events in cardiomyocyte-enriched genes (**Figure S6A-F**). A few exceptions were observed, with metabolically selected iPSC-CMs showing PSI values shifted toward either earlier (**Figure S6G, H**) or later (**Figure S6I, J**) developmental stages relative to antibody-sorted iPSC-CMs.

For iPSC-CM–specific splicing events, both protocols consistently yielded intermediate PSI values for exons that are constitutively spliced in native heart tissue (**Figure 6B–G**), indicating that the splicing differences observed in iPSC-CMs are not attributable to the differentiation method, but rather represent shared features of *in vitro*–derived cardiomyocytes.

For instance, exon 4 of *CLK1* and *CLK4* was more frequently skipped in iPSC-CMs from both protocols compared to heart tissue (**Figure 6B-C**). Exon 4 skipping produces truncated, catalytically inactive isoforms of these kinases ^48^. A trend toward increased exon 4 inclusion was observed during iPSC-CM maturation (**Figure 6B-C**), and significant changes were detected between iPSCs and iPSC-CMs (**Figure 6B-C**).

Increased skipping of *SNRPE* exon 2, which encodes part of the LSM domain required for spliceosome assembly, was also observed in iPSC-CMs relative to heart tissue, regardless of protocol (**Figure 6D**). Unlike *CLK1/4*, this exon was not dynamically regulated during iPSC differentiation or iPSC-CM maturation.

Conversely, *RSRP1* exon 4 showed higher inclusion in iPSC-CMs than in heart tissue, regardless of protocol (**Figure 6E**). Inclusion of this exon disrupts the open reading frame and is predicted to trigger NMD, consistent with lower *RSRP1* expression levels in iPSC-CMs (**Figure S6K**).

Beyond splicing regulators, iPSC-CM-specific patterns were also observed in genes linked to mRNA decay and modification. Inclusion of *UPF3A* exon 4, which encodes a domain required for NMD machinery interactions, was consistently reduced in iPSC-CMs compared to heart tissue (**Figure 6F**). In the m6A methyltransferase *METTL3*, intron 8 retention, which disrupts the methylation domain and produces a truncated, non-functional isoform, was consistently lower in iPSC-CMs (**Figure 6G**). Notably, *METTL3* splicing remained largely unchanged during iPSC differentiation and iPSC-CM maturation (**Figure 6G**).

Finally, we observed differences in contractile gene splicing. In heart tissue, inclusion of *TPM1* exon 2, which encodes part of the tropomyosin domain, exceeded 94%, whereas inclusion was approximately 75% in iPSC-CMs D, G, and T, and ∼88% in iPSC-CMs C25 (**Figure S6L**).

In summary, we identified a subset of unique splicing events that distinguish iPSC-CMs from native cardiac tissue. Despite some protocol-driven variability, most of the splicing patterns were consistent across iPSC-CMs generated with distinct differentiation methods.

Having identified splicing alterations in iPSC-CMs, we next investigated whether changes in the expression of trans-acting splicing regulators could underlie these differences. Our analysis revealed that several mRNAs encoding splicing factors were significantly downregulated in iPSC-CMs compared to pre- and postnatal hearts. These included SR protein kinases CLK1 and SRPK3, as well as ELAVL3, ELAVL4 and NOVA2 (**Table S11**). Cross-referencing with single-cell transcriptomic data from the fetal heart atlas ^23^ showed that most of these genes are predominantly expressed in non-cardiomyocyte cell types. However, *SRPK3* is specifically expressed in cardiomyocytes (**Table S7**), raising the possibility that its reduced expression in iPSC-CMs (**Figure S6M**) may contribute to the altered splicing landscape observed *in vitro*.

## DISCUSSION

In this study, we present a comprehensive characterization of AS dynamics during cardiomyocyte development *in vivo* and *in vitro*, and we highlight discrepancies in splicing regulation between iPSC-CMs and human hearts.

During pre- and postnatal development of the heart, cardiomyocytes undergo a maturation process that involves a wide spectrum of changes in cell structure, metabolism, function and gene expression ^49^. Our comparative analysis of gene expression profiles showed that iPSC-CMs cluster more closely with prenatal heart tissue than postnatal hearts, in agreement with previous reports ^50,51^.

AS is a critical post-transcriptional mechanism that regulates gene expression, affecting over 90% of human genes and allowing a single locus to generate multiple transcript variants with diverse properties ^6,7^. In the heart, developmentally regulated AS has been implicated in the extensive remodeling required to accommodate the increased workload during the transition from fetal to neonatal life and continuing into adulthood ^52–54^. One well-characterized example is the embryonic-specific EH-myomesin isoform, generated by inclusion of *MYOM1* exon 18. This exon encodes a protein domain that enhances elasticity and modulates the viscoelastic proprieties of the sarcomere ^55^. Development is accompanied by a progressive decrease in exon 18 inclusion, and in adult hearts this exon is completely absent (**Fig. S6C**).

Despite recent progress, a comprehensive understanding of splicing regulation throughout human heart development remains incomplete. In our analysis, we identified the majority of alternative splicing events previously well-characterized as regulated during cardiac development *in vivo*. Furthermore, we show that splicing isoforms reported in the developing murine heart are conserved in the human heart (**Table S4**). Most important, we discovered novel splicing events that are differentially regulated between pre- and postnatal human hearts (**Table S4**). While some of these events have been described in the context of disease, their regulation during normal heart development was previously unknown.

In addition to splicing differences between pre- and postnatal hearts, we identified novel splicing events that undergo regulation during prenatal life, between embryonic and fetal stages (**Tables S5 and S6**).

Taken together, our findings offer a detailed perspective on the temporal dynamics of splicing switches during human heart development, uncovering previously uncharted splicing programs associated with cardiac maturation. Importantly, our analysis shows that iPSC-CMs do not correspond to prenatal hearts of a specific gestational age based on their splicing profiles. Instead, splicing patterns in iPSC-CMs are heterogeneous: some events resemble those of early embryonic hearts (**Fig. 4C-F**), others align more closely with later fetal stages (**Fig. 4G-J**), and still a few others mirror the patterns observed shortly after birth (**Fig. S6G**). This variability highlights the heterogeneous maturation state of iPSC-CMs and reflects their incomplete recapitulation of *in vivo* cardiac development.

Additionally, we identified a subset of splicing events that are mis-regulated in iPSC-CMs, with inclusion levels deviating from the constitutive patterns seen *in vivo* (**Tables S9 and S10**). Notably, these iPSC-CM-specific splicing alterations include the mis-splicing of splicing regulators as well as factors involved in mRNA decay and modification. These combined splicing defects suggest widespread disruptions in RNA homeostasis, which may contribute to the functional immaturity of iPSC-CMs compared to native heart cells.

Multiple splicing factors have been implicated in the regulation of cardiac-specific splicing programs. Namely, CELF and MBNL proteins are responsible for many splicing transitions that occur during postnatal heart development in mice ^15,56^, while SRSF1 is critical role for maintaining splicing patterns during postnatal cardiac remodeling ^57^. Additional RNA-binding proteins involved in cardiac splicing regulation include *PTBP1* ^58,59^, *RBFOX2* ^60^, *SRSF5* ^61^, *SRSF10* ^62^, *RBM24* ^63,64^, *RBM20* ^65,66^ and *QKI* ^67^. None of the transcripts encoding these proteins were differentially expressed in iPSC-CMs compared to heart tissue (**Table S11**). However, we found significantly reduced expression of *CLK1* and *SRPK3* in iPSC-CMs (**Table S11**), both of which encode kinases that phosphorylate the serine/arginine-rich (SR) domains of splicing factors ^48,68^. *SRPK3* is of particular interest because it is specifically expressed in cardiomyocytes and phosphorylates RBM20, a key cardiac splicing regulator ^69^. Given that RBM20 activity and nuclear localization depend on its phosphorylation status ^70^, reduced *SRPK3* expression in iPSC-CMs may impair RBM20 function, thus contributing to the altered splicing landscape observed *in vitro*. Importantly, such phosphorylation defects could also help explain the persistence of fetal-like isoforms in iPSC-CMs, as several splicing factors critical for adult splicing programs may remain in a functionally inactive state.

In contrast to *SRPK3*, *CLK1*, another kinase known to phosphorylate RBM20 ^71^, is ubiquitously expressed in the heart. Its downregulation in iPSC-CMs may reflect the absence of non-cardiomyocyte cell types. However, we also observed skipping of the exon encoding the *CLK1* kinase domain in iPSC-CMs, a change likely to impair its enzymatic function. Taken together, these findings support a model in which defective phosphorylation driven by reduced *SRPK3* expression and altered *CLK1* splicing acts upstream of the splicing differences observed in iPSC-CMs.

In conclusion, our findings highlight the ability of iPSC-CMs to model key aspects of early heart development, while also revealing their limitations in recapitulating the full spectrum of splicing programs that govern cardiac maturation. The observed splicing immaturity and variability in iPSC-CMs underscore the need for further optimization of maturation protocols to more faithfully replicate native cardiac splicing transitions, ultimately enhancing their utility as robust *in vitro* models for cardiovascular research.

## Methods

### Human iPSCs

In this study, the following human iPSC lines were used. The DF6.9.9 T.B cell line, provided by the WiCell Bank (https://www.wicell.org/) was reprogrammed from male foreskin fibroblasts ^72^, and is here referred to as iPSC-D. The Gibco® Human Episomal iPSC line (iPSC6.2), here referred to as iPSC-G, was derived from female CD34+ cord blood cells ^73^. The Cuso-2 iPSC line, here referred as iPSC-C, was reprogrammed from skin fibroblasts of a healthy male donor ^74^. The cell line F002.1A.13, here referred to as iPSC-T, was derived from skin fibroblasts of a healthy female donor ^75^. The hiPSC C25 line was a gift from A. Moretti, Munich, Germany ^76^.

### Cardiomyocyte differentiation

Cardiomyocytes differentiated from iPSC lines D, G, T and C were generated as previously described ^77^. On day 13 of differentiation, cellular aggregates were dissociated using 0,25% Trypsin-EDTA (Gibco) for 7 min at 37°C. After dissociation, cells were washed with 2% fetal bovine serum (FBS) and 2mM EDTA in phosphate buffered saline (PBS, 0.1M), and incubated with VCAM1 antibody (BioLegend, 1:50) diluted in PBS/2% FBS for 15 min at 4°C. Then, cells were washed and suspended in PBS/2% FBS for FACS sorting (BD FACSAria™ III Cell Sorter) using a 100μm nozzle at 4°C. Purified VCAM1 positive cells were plated on wells coated with Matrigel, at a seeding density between 20,000 – 40,000 cells/cm2. Alternatively, cardiomyocytes were differentiated from the iPSC C25 line, as described ^78,79^.

### iPSC-CM culture on a micropatterned surface

On day 30 of differentiation, iPSC-CMs were replated by incubating with 0,25% Trypsin-EDTA (Gibco) for 3 min at 37°C, followed by trypsin neutralization using PBS supplemented with 10% FBS. Cells were cultured on micropatterned coverslips (4Dcell, 10mmx10mm, 100 µm pattern) for an additional 10 days. The RPMI + B27 medium was replaced every 2 days and at day 40 of differentiation coverslips were fixed in 3.7% PFA and iPSC-CMs were collected in NZYol (NZYTech®) for subsequent RNA analysis.

### Fluorescence microscopy and image analysis

Immunofluorescence was performed as previously described ^77^ using a mouse anti-myosin-binding protein C antibody (Santa Cruz Biotechnology, sc-131781), a rabbit anti-α-actinin antibody (Abcam, ab9465) and Alexa Fluor™ 546 phalloidin for actin staining. Fluorescence images were acquired with Zeiss LSM 710 Confocal Laser Point-Scanning Microscope. Quantitative image analysis was performed using ImageJ (https://imagej.nih.gov/) standard plugins. For sarcomere length measurements, statistical analysis was performed using Brown-Forsythe and Welch ANOVA tests to account for unequal variances, followed by Games-Howell’s multiple comparisons test.

### Quantitative RT-PCR (qRT-PCR)

Total RNA was extracted using NZYol (NZYTech®) and digested with DNase I (Roche®). cDNA synthesis was performed the SuperScript IV Reverse Transcriptase (Invitrogen^TM^) with oligodT primers. qRT-PCR was carried out using the Universal SYBR Green Supermix (Bio-Rad) and specific primers designed for the selected splicing events (listed in **Table S12**). Each PCR reactions was run in triplicate, using the ViiA™7 RT-PCR Systems (Applied BioSystems). Quantification was performed using the 2–ΔCt method. The level of the target exon was normalized to the level of a constitutive exon from the same gene (i.e., an exon that is consistently included in all splicing isoforms transcribed from the gene). For statistical analysis we used Brown-Forsythe and Welch ANOVA tests to account for unequal variances, followed by Games-Howell’s multiple comparisons test.

### Single-cell RNA sequencing

At day 30, differentiated cells were dissociated using 0.25% Trypsin-EDTA (Gibco) for 5 min at 37°C. After stopping the reaction, dissociated cells were resuspended in PBS/0.1% bovine serum albumin (BSA; Life Technologies), filtered through a 100 μm pore filter and stored in ice. Cell viability was assessed by manual counting using Trypan Blue. Viable cell suspensions (1×106 cells/mL) were partitioned into Gel Beads-in-emulsion (GEMs), in order to capture approximately 10000 cells in single cell droplets, using the Chromium Single Cell 3ʹ GEM, Library & Gel Bead Kit (version 3, 10X Genomics, PN-1000092) and Chromium Single Cell B Chip Kit (10X Genomics, PN-1000074). Barcoded cDNA was extracted from the GEMs by Post-GEM RT-cleanup, PCR amplified and subjected to quality control and quantification. Single Cell 3ʹ Gene Expression libraries from single cells was then performed using Chromium i7 Multiplex Kit (PN-120262). All the steps were performed according to the manufacturer’s protocol. Libraries were sequenced using an Illumina NextSeq 500 Instrument with a 300 cycles kit. The raw FASTQ files have been uploaded to ArrayExpress and can be accessed using the accession number E-MTAB-13850.

### Single-cell RNA sequencing analysis

Raw sequencing reads from FASTQ files were pseudo-aligned to the transcriptome using Kallisto BUStools ^80^. The resulting count matrix was imported into R and converted into a Seurat object using Seurat v.5.1.0 ^81^. To retain high-quality cell-containing droplets, emptyDrops from DropletUtils v1.24.0 ^82^ was used with a false discovery rate (FDR) of ≤ 0.01. Cells were filtered according to their number of expressed genes (≥ 500), number of UMIs (≥ 1000), log10GenesPerUMI (> 0.8) and ribosomal ratio (> 0.05). The threshold for the expression of mitochondrial genes was set to 40% given the high energy needs of cardiomyocytes ^83^. We further kept only those genes expressed in more than 10 cells in each sample. Counts were normalized, adjusted for variance, and the 1000 most highly variable features were identified using the SCTransform function, while regressing out sequencing depth effects. Principal component analysis was performed on these features and the five principal components explaining the most variance were used for UMAP dimensionality reduction and to build a SNN graph. Known cardiomyocyte markers were used to select an appropriate resolution that allows to capture this cell population. Clustering was done using FindClusters function, with the Louvain algorithm with multilevel refinement, and resolution set to 0.01 for iPSC-CM_T and 0.05 for iPSC-CM_D. Doublet detection and removal were performed using DoubletFinder v.2.0.4 with the first 5 principal components (PCs) and pK = 0.07 for iPSC-CM_T and pK = 0.05 for iPSC-CM_D. Cluster markers were identified using Wilcoxon’s rank sum test implemented in FindAllMarkers function. We considered only positive markers, as well as genes detected in at least 25% of the cells of each population, and with a minimum log2FC of 0.5. Markers with an adjusted p-value ≤ 0.01 were ranked by log₂FC and used for gene set enrichment analysis (GSEA) with gseGO function of clusterProfiler 4.12.6 ^84^ and org.Hs.eg.db v.3.16.0 ^85^. For automated cell type classification, label transfer was performed using Seurat’s FindTransferAnchors and TransferData functions, using the first 25 PCs of Farah *et al.* dataset ^23^, using the processed Seurat object publicly available. Only cells assignments with a prediction score higher than 0.6 were kept.

### Bulk RNA-sequencing

RNA was extracted using NZYol (NZYTech®) and digested with DNase I (Roche®). At least 1µg of total RNA was obtained from each culture of iPSC-CMs at day 30 of differentiation. After purification with poly-T oligo-attached magnetic beads, libraries were prepared using the NEBNext® Ultra™ Directional RNA Library Prep Kit, and sequenced on the Illumina platform Novaseq6000. The raw FASTQ files have been uploaded to ArrayExpress and can be accessed using the accession number E-MTAB-13757 ^28^.

Additionally, we used RNA sequencing data from 50 healthy human hearts - 38 prenatal and 12 postnatal, generated in a previous study. Raw data (FASTQ files) was downloaded from ArrayExpress (https://www.ebi.ac.uk/arrayexpress/), with the accession number E-MTAB-6814 39.

### Data preprocessing, genome alignment and transcript quantification

Quality control of the sequenced reads was performed using *FastQC v0.11.5* ^86^. Adaptors, low-quality bases (Phred quality <20) and reads shorter than 75bp were filtered out using *Trimgalore v0.6.4* ^87^. Filtered reads were pseudo-aligned to the human transcriptome and bootstrapped transcript abundance was estimated with *Salmon v1.10.2* ^88^, with default parameters, 100 resamplings, fragment length distribution set to a mean of 100 bp and standard deviation of 10 bp, and correction for GC and sequencing bias. Trimmed reads were also mapped to the human reference genome (GRCh38/hg38) using the splice-aware aligner *STAR v2.7.10b* ^89^. Chimeric alignments were detected (--chimSegmentMin 20), splice junctions were optimized with -- sjdbOverhang 99, and two-pass mapping (--twopassMode Basic) was used to improve splice junction detection. Only uniquely mapped reads were kept using *samtools v.1.7* ^90^. GENCODE v45 human primary gene annotation was used as reference ^91^. **Table S1** summarizes the main biological and technical characteristics of all the samples.

### Gene expression analysis

Transcriptomic analysis was performed using R *v4.4.1* software ^92^. Prior to further analysis, data was filtered so that only genes with more than 10 read counts in at least three samples were kept. Transcript read counts were converted into a gene-level count matrix using *tximport v1.32.0* ^93^. To filter out lowly expressed genes that could possibly result from transcriptional noise, only genes with transcripts per million (TPM) > 1 were considered. This threshold was determined based on the read counts distribution observed with a log10(TPM) histogram. Pearson pairwise correlation coefficients (r) were computed on rlog-transformed counts to assess the similarity between samples. The correlation matrix was visualized using *corrplot v0.*94 ^94^. Venn diagrams illustrating the overlap of expressed genes (TPM > 1) between different iPSC-CM lines were generated using *VennDiagram v1.7.3* ^95^. To prevent highly expressed genes from dominating the analysis, counts were transformed using varianceStabilizingTransformation (VST) from *DESeq2 v1.44.0* ^96^. Principal component analysis (PCA) was performed on transformed read counts to discern the main sources of variation in the dataset using *ggplot2 v3.5.1* ^97^. Regular hierarchical clustering dendrograms were drawn using the R base function hclust with average linkage clustering and 1-pairwise Spearman correlation coefficients as the distance metric. The *pvclust v2.2-0* ^98^ package was used to assess the uncertainty of the clustering and to compute p-values for each cluster based on 1000 bootstrap resamplings.

### Analysis of Metabolic Pathways

To evaluate pathway-specific changes in metabolism, we performed single-sample Gene Set Enrichment Analysis (ssGSEA), which enables the estimation of gene set enrichment scores across individual samples. Curated hallmark gene sets for the Glycolysis (MSigDB ID: M5937) and Oxidative Phosphorylation (MSigDB ID: M5936) pathways were obtained from the Molecular Signatures Database (MSigDB) using the *msigdbr* package *v.24.1.0* ^99^. ssGSEA was applied to the VST expression matrix to quantify pathway activity for each sample using the gsva() function ^100^. Enrichment scores for the Glycolysis and OXPHOS pathways were visualized using violin plots to illustrate their distribution across iPSC-CMs, prenatal and postnatal hearts. To complement the pathway-level analysis, we conducted a targeted gene expression analysis focusing on key enzymes and regulators involved in cardiac metabolic processes. This allowed for finer resolution of developmental expression trends across glycolysis, the TCA cycle, oxidative phosphorylation (OXPHOS), and fatty acid oxidation (FAO). Representative genes were manually curated from each pathway based on their central metabolic roles and known developmental relevance in cardiac tissues. A heatmap was generated using the *pheatmap* package to visualize standardized (row-scaled) VST values of the selected genes across all developmental stages.

### Deconvolution of bulk RNA-seq

Cell-type proportions in heart tissue and iPSC-CM bulk RNA-seq samples were estimated using *CIBERSORTx* ^101^. Single-cell RNA-seq data from prenatal hearts (Farah *et al.* single-cell atlas ^23^) was used as reference and converted into a signature matrix using 50 replicates per cell type and retaining only genes with an average expression > 0.5 across cells. To estimate cell-type composition, *CIBERSORTx* Fractions was applied with S-mode batch correction to account for technical variability, and statistical significance was assessed with 1000 permutations for p-value calculation. A heatmap was generated to visualize cell-type proportions where atrial, ventricular, and nodal-like cardiomyocytes were merged into a single category of cardiomyocytes; red blood and white blood cells into hematopoietic; BEC, LEC and endocardial cells into endothelial cells; and fibroblasts, smooth muscle cells and epicardial into mesenchymal cells.

### Expression of genes per cell-type

To explore the expression of selected genes across cardiac cell types, we used the processed Seurat object provided by the authors of Farah et al. (2024), which includes fetal human heart single-cell RNA-seq data spanning developmental stages from 9 to 15 post-conceptional weeks ^23^. Cardiomyocyte subtypes - atrial, ventricular, and nodal-like - were grouped under a single cardiomyocyte category, to facilitate comparisons across major cardiac lineages. A dot plot was generated to summarize expression levels across cell types for selected genes.

### Splicing analysis

Alternative splicing (AS) events were identified and quantified using *rMATS v4.1.2*, *MAJIQ v2.5.1,* and *vast-tools v2.5.1* ^102–104^, which extract reads mapping to splice junctions from BAM files and calculate Percent Spliced In (PSI) values. For *rMATS*, default parameters were used (cstat 0.0001, anchorLength 3) with the -novelSS option enabled to detect unannotated splice sites. Results were filtered to retain events with ≥10 reads supporting either inclusion or skipping in at least half of the samples across at least two groups. For *MAJIQ*, a splice graph was constructed, including all binary splicing events detected in >50% of the samples per group, with a minimum of 5 supporting reads for known junctions and 7 reads for *de novo* junctions. For *vast-tools*, PSI values were computed, including intron retention events (--p_IR). Events were retained if they had sufficient read coverage in 50% of both prenatal and postnatal groups (--min_Fr 0.5), a minimum standard deviation of 5 (min_SD 5), and met the --noVLOW criterion to exclude low-coverage events. All splicing events were further filtered to retain only those with ≤1/3 missing values in prenatal and postnatal heart samples and no missing values in iPSC-CM samples.

To assess splicing similarity between samples, Spearman pairwise correlation coefficients (ρ) were computed on logit-transformed PSI values. To determine a TPM threshold for reliable PSI estimation, splicing events were binned based on the TPM of the parent gene, and the mean PSI variance of each bin was plotted. A loess smooth curve was fitted to identify the stabilization point, determining TPM > 10 as optimal for robust splicing estimation. All splicing events detected by rMATS in genes with TPM > 10 were considered, except for constitutive events where all samples exhibited a PSI of 0 or 1. The correlation matrix was generated and visualized using *corrplot v0.94*^94^. PCA was performed using PSI values from the 500 most variable splicing events and visualized using *ggplot2 v3.5.1* ^97^.

Significant splicing changes between groups were defined as those events with deltaPSI > +/- 0.2 and < 1% FDR for *rMATS*. With *MAJIQ* heterogen and modulize, we considered as significant binary events with a deltaPSI > +/- 0.2 and a probability of changing > 0.9. *Vast-tools* was also used to detect differentially spliced events with > 90% probability that there is a change of at least 20%.

The heatmap to visualize the differentially spliced events was generated using *pheatmap v1.0.12* ^105^ and *RColorBrewer v.1.1* ^106^. For this, each sample’s deviation from the mean of each gene was calculated as row Z-scores, and clustering was performed using complete linkage clustering with Spearman correlation as the distance metric. Visualization of specific alternative splicing events was performed with *ggashimi v1.1.5* using the respective surrounding coordinates and the median number of supporting reads for each group ^107^. The potential functional impact of the alternative splicing events was investigated using *VastDB*, by matching the event coordinates with those present in the database ^104^. Unless specified otherwise, all other plots were generated using the *ggplot2 v3.5.1* package ^97^. When multiple transcript isoforms corresponded to a specific spliced exon/intron, we reported the isoform with the highest confidence (i.e., MANE Select transcript, ENSEMBL golden isoform, or similar high-confidence annotations), based on the GENCODE v45 human primary transcript annotation used throughout the analysis ^91^.

### NEASE Analysis

*NEASE* (Network-based Enrichment Analysis for Splicing Events) *v1.2.2* was used to infer the functional impact of the identified AS events at the protein interaction network level ^108^. Exons predicted to disrupt the open reading frame (ORF) were excluded from the analysis. First, we detected protein features potentially impacted by splicing, including protein domains, linear motifs and interacting residues. Then, pathways affected by interactions mediated by the spliced features were identified. One-sided hypergeometric test was performed to identify enriched pathways, applying a p-value cutoff of 0.05. Pathways were ranked based on the NEASE score.

### Sample preparation for MS analysis

Cardiomyocytes differentiated from iPSC lines D, T and C were collected at day 30 through centrifugation and the resulting pellet was stored at −80 °C. Cell pellets were dissolved in lysis buffer containing 5% sodium dodecyl sulfate (SDS) and 50 mM triethylammonium bicarbonate (TEAB), pH 8.5, and each sample was transferred to a PIXUL^TM^ 96-well plate (Active Motif). Samples were sonicated with a PIXUL^TM^ Multisample sonicator (Active Motif) for 5 minutes with default settings. After centrifugation for 15 min at 2204 x g at room temperature (RT), the lysates were transferred to a new plate and the protein concentration was measured by bicinchoninic acid (BCA) assay (Thermo Scientific). From each sample 50 µg of protein was isolated and volumes were adjusted to 100 µl with the lysis buffer to continue the protocol. Proteins were reduced by addition of 15 mM dithiothreitol and incubation for 30 minutes at 55°C and then alkylated by addition of 30 mM iodoacetamide and incubation for 15 minutes at RT in the dark. Phosphoric acid was added to a final concentration of 1.2% and subsequently samples were diluted 7-fold with binding buffer containing 90% methanol in 100 mM TEAB, pH 7.55. The samples were loaded on a 96-well S-Trap^TM^ plate (Protifi) in parts of 400 µl, placed on top of a deepwell plate, and centrifuged for 2 min at 1,500 x g at RT. After protein binding, the S-trap^TM^ plate was washed three times by adding 200 µl binding buffer and centrifugation for 2 min at 1,500 x g at RT. A new deepwell receiver plate was placed below the 96-well S-Trap^TM^ plate and 125 µl 50 mM TEAB containing 1 µg trypsin (1/50, w/w) was added for digestion overnight at 37°C. Using centrifugation for 2 min at 1,500 x g, peptides were eluted in three times, first with 80 µl 50 mM TEAB, then with 80 µl 0.2% formic acid (FA) in water and finally with 80 µl 0.2% FA in water/acetonitrile (ACN) (50/50, v/v). Eluted peptides were dried completely by vacuum centrifugation.

TMTpro^TM^ 16-plex labels (0.5 mg, lot #WF322713, Thermo Fisher Scientific) were equilibrated to RT immediately before use and dissolved in 20 µl anhydrous acetonitrile (ACN). The dried peptides were re-suspended in 90 µl 100 mM TEAB (pH 8.5), peptide concentration was determined on a Lunatic spectrophotometer (Unchained Labs) ^109^ and peptide amount was adjusted to 35 µg for each sample. Peptides were labeled for 1 hour at RT using 0.25 mg of TMTPro^TM^ label (127N – C; 128C – T; 133N – D). The reaction was quenched for 15 min at RT by addition of 4.2 µl 5% hydroxylamine. The labeled samples were combined, 100 µg labeled peptides was isolated, dried by vacuum centrifugation, re-dissolved in 100 µl loading solvent A (0.1% TFA in water/ACN (98:2, v/v)), pH was adjusted with 1 µl of 100% TFA and the sample was desalted on a reversed phase (RP) C18 OMIX tip (Agilent). The tip was first washed 3 times with 100 µl pre-wash buffer (0.1% TFA in water/ ACN (20:80, v/v)) and pre-equilibrated 5 times with 100 µl of wash buffer (0.1% TFA in water) before the sample was loaded on the tip. After peptide binding, the tip was washed 3 times with 100 µl of wash buffer and peptides were eluted twice with 100 µl elution buffer (0.1% TFA in water/ACN (40:60, v/v)). The combined elutions were dried in a vacuum concentrator.

Vacuum dried peptides were re-dissolved in 100 µl loading solvent A and 95 µl (+/- 100 µg) was injected for fractionation by RP-HPLC (Agilent series 1200) connected to a Probot fractionator (LC Packings). Peptides were first loaded in loading solvent A on a 4 cm pre-column (made in-house, 250 µm internal diameter (ID), 5 µm C18 beads, Dr. Maisch) for 10 min at 25 µl/min and then separated on a 15 cm analytical column (made in-house, 250 µm ID, 3 µm C18 beads, Dr Maisch). Elution was done using a linear gradient from 100% RP-HPLC solvent A (10 mM ammonium acetate (pH 5.5) in water/ACN (98:2, v/v)) to 100% RP-HPLC solvent B (70% ACN, 10 mM ammonium acetate (pH 5.5)) in 100 min at a constant flow rate of 3 µL/min. Fractions were collected every minute between 20 and 92 minutes and pooled every 24 minutes to generate a total of 24 samples for LC-MS/MS analysis. All 24 fractions were dried under vacuum in HPLC inserts and stored at −20°C until further use.

### Liquid chromatography-MS/MS analysis

Purified peptides were re-dissolved in 20 µl solvent A and 15 µl of each fraction was injected for LC-MS/MS analysis on an Ultimate 3000 RSLCnano system in-line connected to an Orbitrap Fusion Lumos mass spectrometer equipped with a pneu-Nimbus dual ion source (Phoenix S&T). Trapping was performed at 20 μl/min for 2 min in solvent A on a 5 mm PepMap™ Neo trapping column (300 μm ID, 5 μm beads, C18, Thermo) and the sample was loaded on a 110 cm prototype µPAC column (Thermo Scientific™) with C18-endcapped functionality mounted in the Ultimate 3000’s column oven at 50°C. For proper ionization, a fused silica PicoTip emitter (10 µm inner diameter) (New Objective) was connected to the µPAC™ outlet union and a grounded connection was provided to this union. Peptides were eluted by a non-linear increase from 2 to 26.4% MS solvent B (0.1% FA in water/ACN (2:8, v/v)) over 45 minutes, then to 44% in 55 minutes, first at a flow rate of 600 nl/min, then at 300 nl/min, followed by a 5-minutes wash at 60 minutes reaching 56% MS solvent B and re-equilibration with MS solvent A (0.1% FA in water).

The mass spectrometer was operated in data-dependent mode (TMT pro SPS on Lumos) with a top speed of three seconds. Full-scan MS spectra (375-1500 m/z) were acquired at a resolution of 120,000 in the Orbitrap analyzer after accumulation to a target AGC value of 400,000 with a maximum injection time of 50 ms. The precursor ions were filtered for charge states (2-7 required), dynamic exclusion (60 s; +/- 10 ppm window) and intensity (minimal intensity of 5E4). The precursor ions were selected in the quadrupole with an isolation window of 0.7 Da and accumulated to an AGC target of 1E4 or a maximum injection time of 50 ms and activated using CID fragmentation (35% NCE). The fragments were analyzed in the Ion Trap Analyzer at turbo scan rate. The 10 most intense MS2 fragments were selected in the quadrupole using MS3 multi-notch isolation windows of 2 m/z. An orbitrap resolution of 60,000 was used with an AGC target of 1E5 or a maximum injection time of 118 ms and activated using HCD fragmentation (65% NCE). The polydimethylcyclosiloxane background ion at 445.120028 Da was used for internal calibration (lock mass). QCloud was used to control instrument longitudinal performance ^110^.

### MS data analysis

LC-MS/MS data from the iPSC-CMs generated in this study (PRIDE accession: PXD000000) and from the publicly available dataset PXD006675 ^47^, which includes adult heart samples (ventricle, auricle, and septum), were analyzed using MaxQuant software (version 2.6.4.0). Searches were performed using default parameters, including a false discovery rate (FDR) of 1% at both the peptide and protein levels. Correction factors were used and extracted from TMT correction factor certificate from manufacturing lot #WF322713.

Protein identification was carried out using FASTA reference files from the UniProt human proteome (UP000005640_9606 and UP000005640_9606_additional; release 2025_04). Enzyme specificity was set to cleavage C-terminal to arginine and lysine residues, including cleavage at Arg/Lys–Pro bonds, allowing up to two missed cleavages. Carbamidomethylation of cysteine was set as a fixed modification, while methionine oxidation and N-terminal acetylation were included as variable modifications. The “match between runs” feature was enabled to enhance peptide identification across LC-MS/MS runs. Peptide identification for selected genes was extracted from the “peptides.txt” output file.

## Acknowledgements

We are grateful to Leslie Leinwand (University of Colorado, Boulder, CO, USA) and Lars Steinmetz (Stanford University, Palo Alto, CA, USA) for insightful discussions. We also thank the GIMM Flow Cytometry, Bioimaging and Genomics Facilities (Lisboa, Portugal), the BIH/MDC Genomics Technology Platform (Berlin, Germany), and Carmen Judis (MDC, Berlin, Germany) for technical support. Mass spectrometry-based proteomics services were provided by the VIB Proteomics Core (Ghent, Belgium), whose support is gratefully acknowledged.

## Author Contributions

B.G.-S., M.F. and M.C.-F designed the study. B.G.-S. performed the bioinformatics analyses. M.F., M.R., S.M., M.T.C. performed the experiments involving iPSC-CM differentiation and characterization. A.V.-G. performed the proteomic analysis. P.N.P. contributed RNA-seq data from iPSC-CM line C25. H.M. contributed to scRNA-sequencing. R.S. and M.G. provided expert guidance and feedback on analyses and results. B.G.-S., M.F. and M.C.-F wrote the manuscript, with feedback from all authors.

---

**Supplementary Figure 1.**
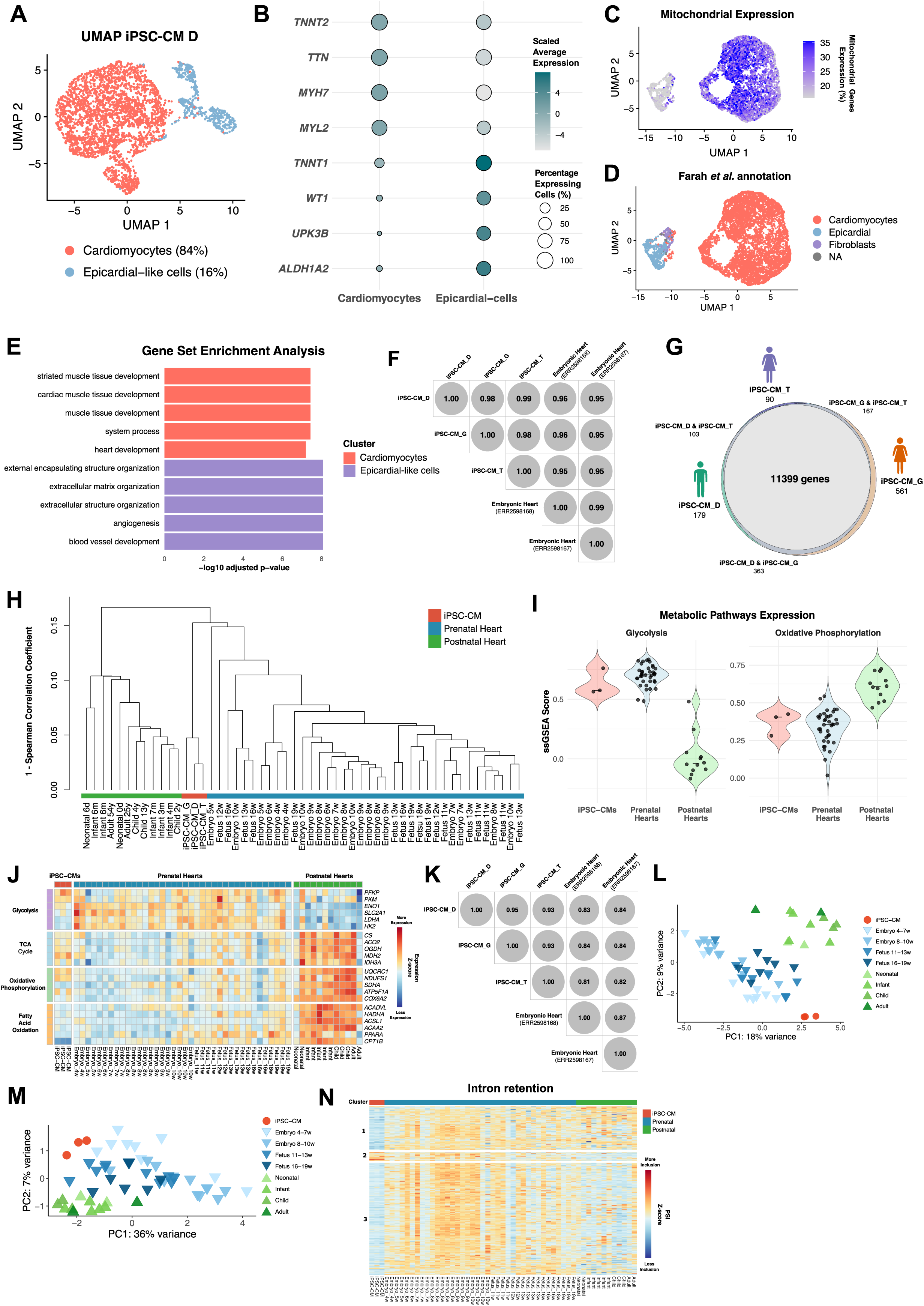
Genome-wide comparison of splicing profiles in iPSC-CMs and developing hearts. **A.** UMAP plot of iPSC-CMs derived from iPSC-D, revealing two major subpopulations of cells corresponding to cardiomyocytes (red), and epicardial-derived cells (blue). **B.** Dot plot of the expression of markers for the cell types identified in (**A**). Dot size represents the percentage of cells expressing each gene, and dot color indicates scaled expression levels averaged per cell population. **C.** UMAP plots displaying the enriched expression of mitochondrial genes in iPSC-CM derived from iPSC-T. **D.** UMAP plot of iPSC-CMs derived from iPSC-T with cell type annotations based on label transfer from the Farah *et al.* single-cell atlas. **E.** Gene set enrichment analysis of population-specific markers with adjusted p-value <= 0.01 sorted by average log2FC. **F.** Correlation matrix showing gene expression similarities (TPM > 1) between iPSC-CM cultures derived from iPSC-D, iPSC-G, and iPSC-T, and two distinct 4-week embryonic hearts. Gene expression counts were rlog-transformed, and pairwise correlation coefficients were computed using Pearson correlation. **G.** Venn diagram illustrating the overlap of the protein-coding genes expressed (TPM>1) in iPSC-CM cultures derived from iPSC-D (green), iPSC-G (orange) and iPSC-T (purple). **H.** Hierarchical clustering of all samples, based on 1 – pairwise Spearman correlation coefficients, utilizing the average linkage method. Samples are colored according to their group (iPSC-CMs - red; prenatal heart – blue; postnatal heart - green). **I.** Violin plots showing the single-sample GSEA score of iPSC-CMs, prenatal and postnatal hearts for Glycolysis and Oxidative Phosphorylation (OXPHOS) pathways. **J.** Heatmap displaying the Z-score normalized expression levels of key genes involved in (from top to bottom) glycolysis, the TCA cycle, oxidative phosphorylation (OXPHOS) and fatty acid ß-oxidation (FAO), across iPSC-CMs, prenatal and postnatal hearts. **K.** Correlation matrix showing the inclusion levels (TPM>10) of all alternative splicing events detected by rMATS in iPSC-CM cultures derived from iPSC-D, iPSC-G and iPSC-T. Constitutive events with a Percent Spliced In (PSI) of 0 or 1 were removed. PSIs were logit-transformed and pairwise correlation coefficients were calculated using Spearman correlation. **L-M.** PCA of the 500 most variable splicing events in iPSC-CMs (red), prenatal hearts (in blue) and postnatal hearts (in green), as identified by (**L**) MAJIQ and (**M**) rMATS. Lighter colors represent younger samples, while darker colors represent older samples. **N.** Heatmap displaying Z-score normalized Percent Spliced In (PSI) levels for all intron retention events, comparing iPSC-CMs with prenatal hearts, iPSC-CMs with postnatal hearts, and prenatal hearts with postnatal hearts. Colors reflect inclusion relative to the mean of each event (red for higher inclusion, blue for lower inclusion). Event-specific information is provided in Tables S2 and S3.

**Supplementary Figure 2.**
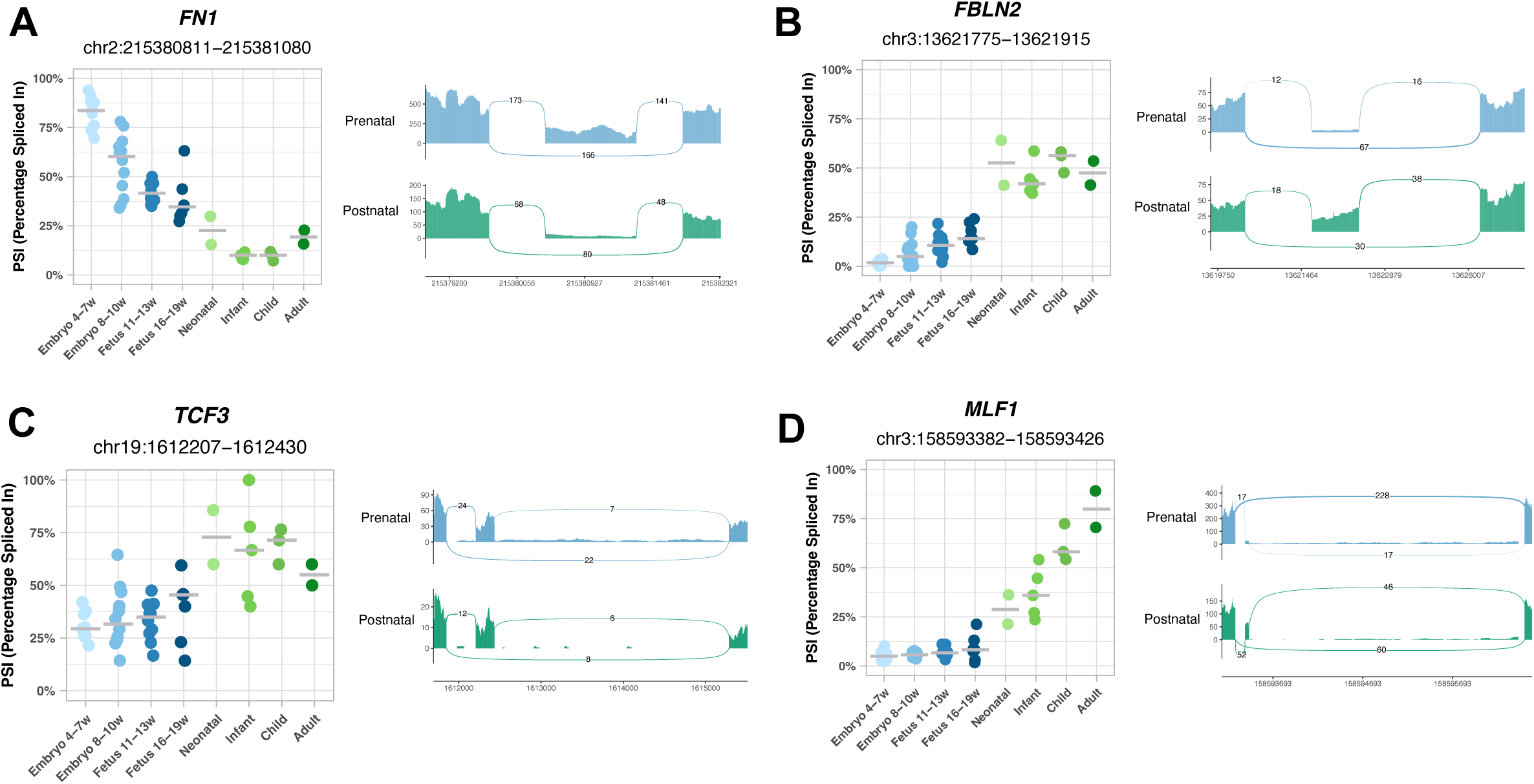
Pre-to postnatal splicing transitions in the heart. **A-D.** Inclusion levels (PSI) of exons differentially included between prenatal and postnatal hearts, specifically (**A**) *FN1* exon 33 (*FN1*-203), (**B**) *FBLN2* exon 9 (*FBLN2*-203), (**C**) *TCF3* exon 18 (*TCF3*-203), (**D**) *MLF1* exon 3 (*MLF1*-203). PSI levels are shown for prenatal and postnatal hearts, across various developmental time points. Each event is accompanied by the corresponding sashimi plot.

**Supplementary Figure 3.**
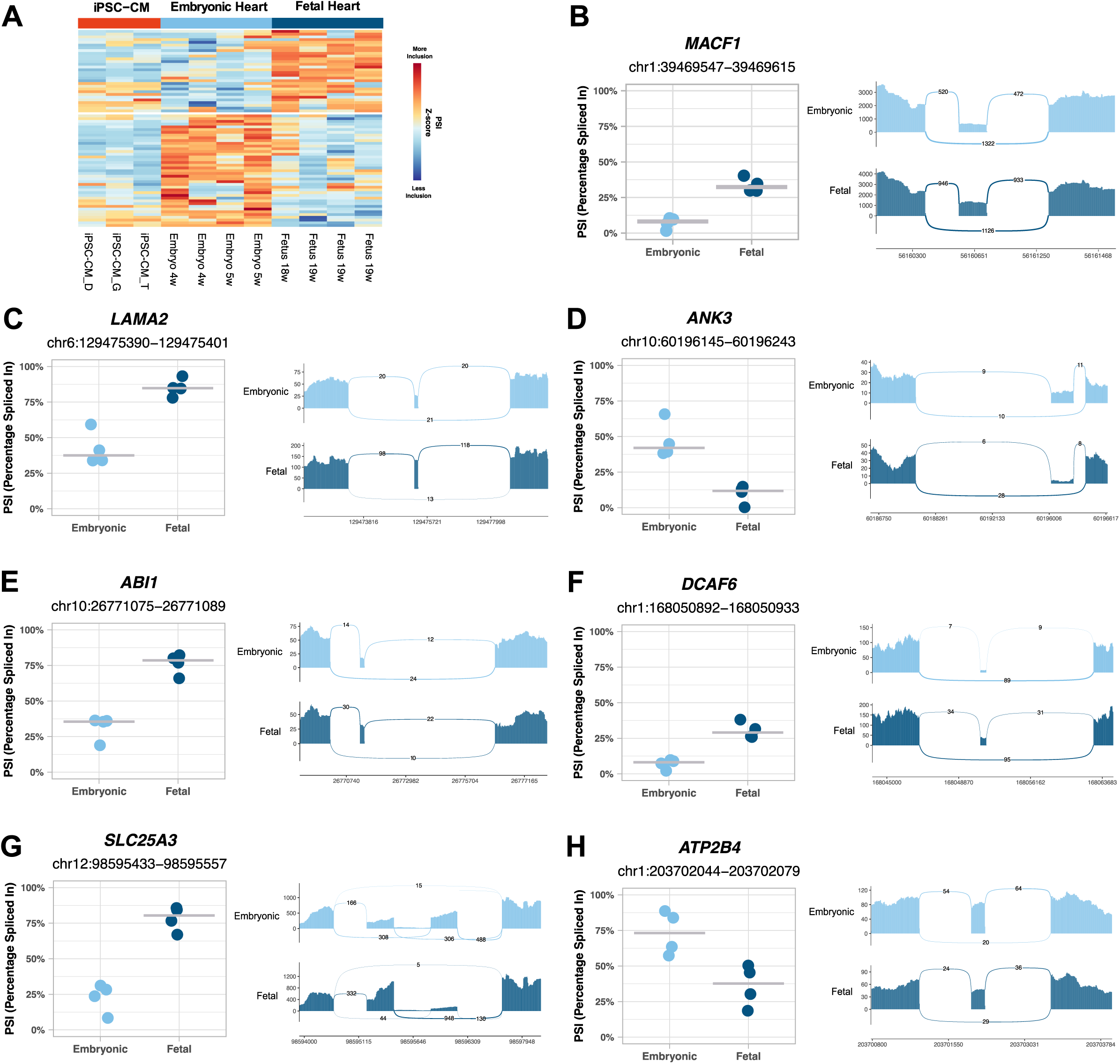
Splicing regulation in the developing prenatal hearts. **A.** Heatmap displaying Z-score normalized Percent Spliced In (PSI) levels of all intron retention found when comparing 4-5 weeks embryonic with 18-19 weeks fetal hearts. Colors represent inclusion levels relative to the mean for each event (red indicates higher inclusion, blue indicates lower inclusion). **B-H.** Inclusion levels (PSI) of exons differentially spliced between 4-5 weeks embryonic and 18-19 weeks fetal hearts, specifically (**B**) *MYO18A* exon 11 (*MYO18A*-203), (**C**) *LAMA2* exon 53 (*LAMA2*-201), (**D**) *ANK3* exon 16 (*ANK3*-201), (**E**) *ABI1* exon 4 (*ABI1*-206), (**F**) *DCAF6* exon 17 (*DCAF6*-202), (**G**) *SLC25A3* exon 3 (*SLC25A3*-202) and (**H**) *ATP2B4* exon 7 (*ATP2B4*-203). PSI levels are shown for embryonic hearts and fetal hearts. Each event in accompanied the corresponding sashimi plot.

**Supplementary Figure 4.**
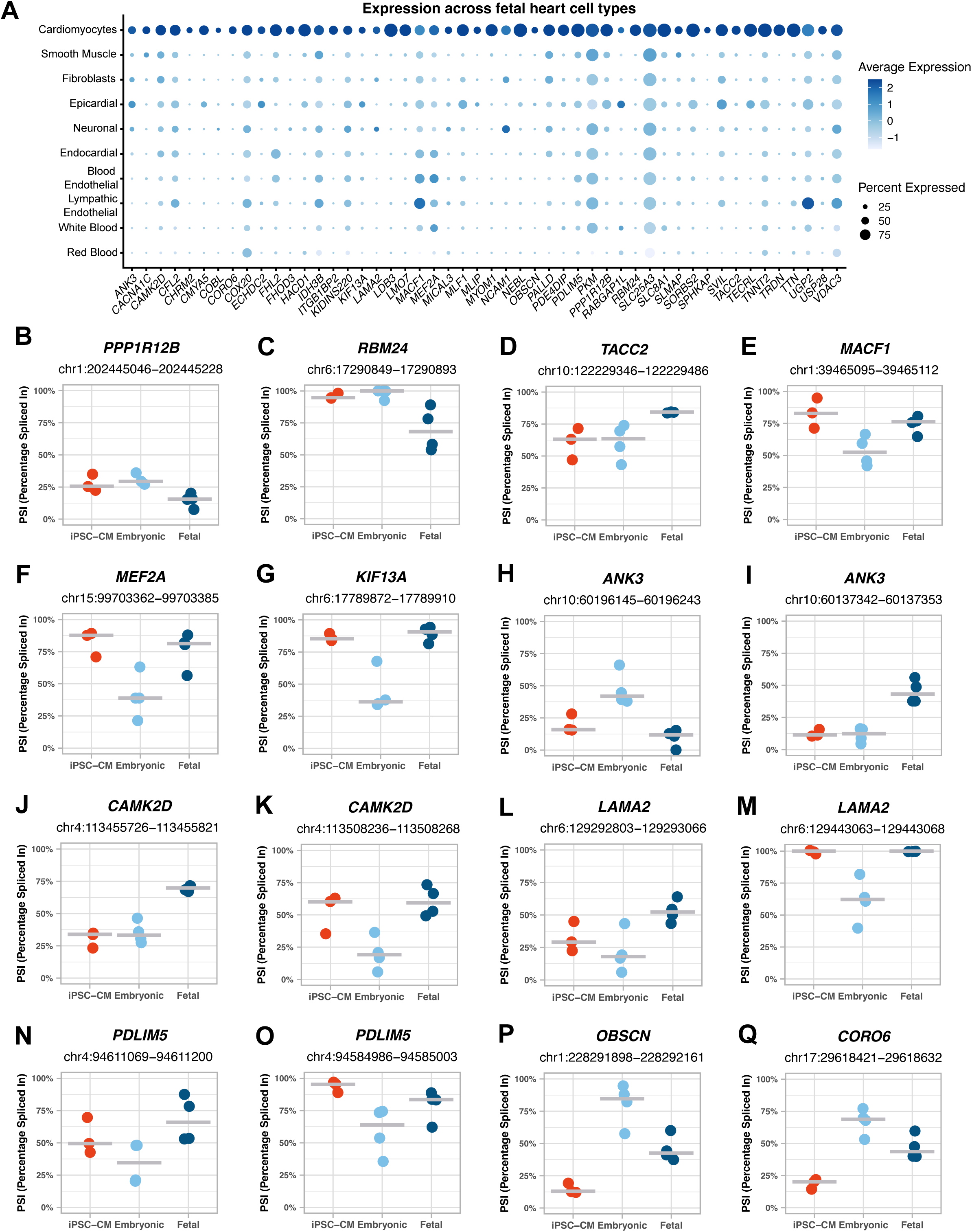
Heterogeneous mirroring of prenatal heart splicing regulation in iPSC-CMs. **A.** Dot plot depicting the expression of cardiomyocyte-specific genes (log2FC > 0.5 compared with all other cell types) across the fetal heart single-cell atlas. Dot size represents the percentage of cells expressing each gene, and dot color indicates the scaled expression levels averaged per cell population. **B-Q**. Inclusion levels (PSI) of exons differentially spliced between 4-5 weeks embryonic and 18-19 weeks fetal hearts, specifically **(B)** *PPP1R12B* exon 13 (*PPP1R12B*-204), (**C**) *RBM24* exon 4 (*RBM24*-207), **(D)** *TACC2* exon 15 (*TACC2*-210), (**E**) *MACF1* exon 95 (*MACF1*-230), (**F**) *MEF2A* exon 9 (*MEF2A*-205), (**G**) *KIF13A* exon 26 (*KIF13A*-201), (**H**) *ANK3* exon 16 (*ANK3*-201), (I) *ANK3* exon 25 (*ANK3*-209), (**J**) *CAMK2D* exon 20 (*CAMK2D*-211), (**K**) *CAMK2D* exon 14 (*CAMK2D*-237), (**L**) *LAMA2* exon 21 (*LAMA2*-206), (**M**) *LAMA2* exon 44 (*LAMA2*-201), (**N**) *PDLIM5* exon 11 (*PDLIM5*-222), (**O**) *PDLIM5* exon 6 (*PDLIM5*-207), (**P**) *OBSCN* exon 44 (*OBSCN*-228), (**Q**) *CORO6* exon 4 (*CORO6*-216). PSI levels are shown for iPSC-CMs, embryonic hearts and fetal hearts.

**Supplementary Figure 5.**
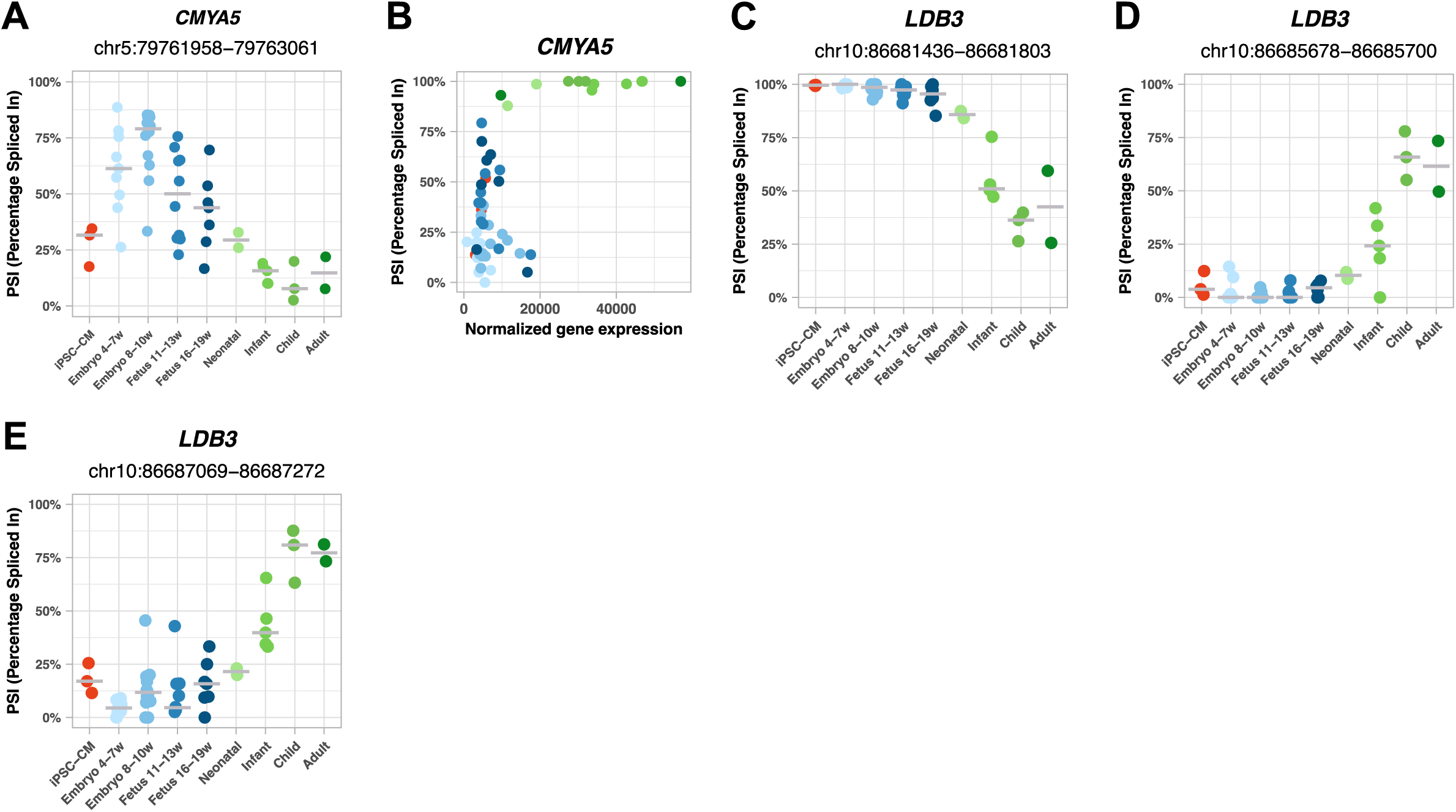
Maturation-driven splicing changes in iPSC-CMs recapitulate developmental transitions in the human heart. **A, C-E**. Inclusion levels (PSI) of exons/introns differentially spliced between prenatal and postnatal hearts, specifically (**A**) *CMYA5* intron 8 (*CMYA5*-201), (**C**) *LDB3* exon 5 (*LDB3*-202), (**D**) *LDB3* exon 5 (*LDB3*-205) and (**E**) *LDB3* exon 6 (*LDB3*-205). PSI levels are shown for iPSC-CMs, prenatal hearts, and postnatal hearts across various developmental time points. **B.** Correlation between the Percent Spliced In of *CMYA5* exon 5, which is predicted to lead to NMD when spliced out, and *CMYA5* gene expression levels.

**Supplementary Figure 6.**
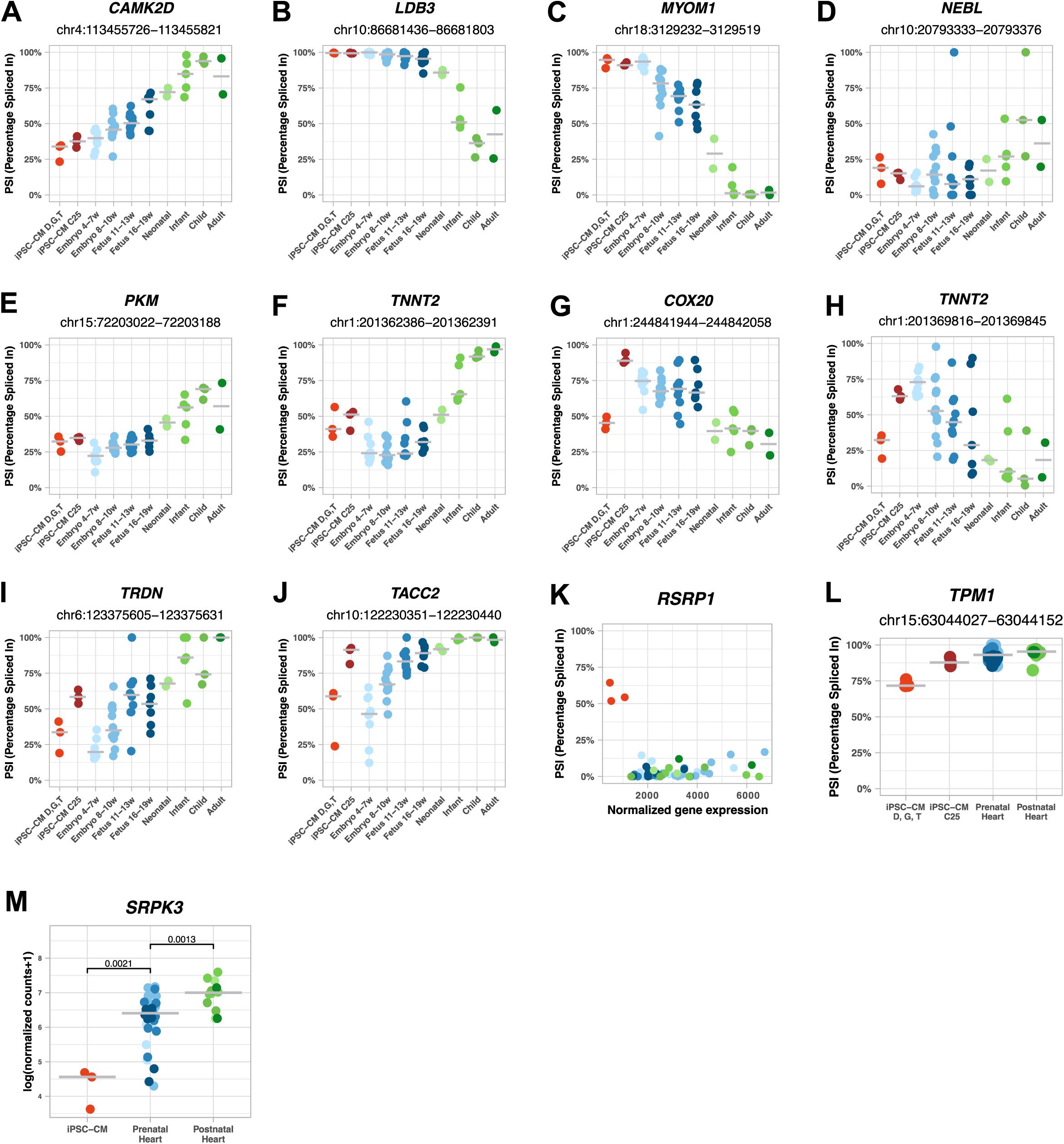
Protocol-independent iPSC-CM–specific splicing events. **A-J**. Inclusion levels (PSI) of exons in iPSC-CMs and both prenatal and postnatal hearts, specifically (**A**) *CAMK2D* exon 20 (*CAMK2D*-211), (**B**) *LDB3* exon 5 (*LDB3*-205), (**C**) *MYOM1* exon 18 (*MYOM1*-202), (**D**) *NEBL* exon 6 (*NEBL*-224), (**E**) *PKM* exon 9 (*PKM*-219), (**F**) *TNNT2* exon 10 (*TNNT2*-203), (**G**) *COX20* exon 2 (*COX20*-203), (**H**) *TNNT2* exon 8 ( *TNNT2*-202), (**I**) *TRDN* exon 19 (*TRDN*-201), (**J**) *TACC2* exon 16 (*TACC2*-210). PSI levels are shown for iPSC-CMs purified by fluorescence-activated cell sorting with VCAM1 antibodies (iPSC-CMs D, G, T), iPSC-CMs purified by metabolic selection with lactate (iPSC-CM C25), prenatal hearts, and postnatal hearts. **K.** Correlation between the Percent Spliced In of *RSRP1* exon 4, which is predicted to lead to NMD when included, and *RSRP1* gene expression levels (DESeq2 normalized counts). **L.** Inclusion levels (PSI) of *TPM1* exon 2 (*TPM1*-207) across iPSC-CMs purified by fluorescence-activated cell sorting with VCAM1 antibodies (iPSC-CMs D, G, T), iPSC-CMs purified by metabolic selection with lactate (iPSC-CM C25), prenatal and postnatal hearts. **M.** Normalized expression of *SRPK3* in iPSC-CMs, prenatal hearts, and postnatal hearts.

## Supplemental Information

**Document S1**. Figures S1-S6

**Table S1**. Sequencing metrics of single-cell and bulk RNA-sequencing from iPSC-CMs.

**Table S2**. Differentially spliced events detected between prenatal and postnatal hearts.

**Table S3**. Differentially spliced events detected between iPSC-CMs and prenatal hearts, and between iPSC-CMs and postnatal hearts.

**Table S4**. Selected AS events between prenatal and postnatal hearts, along with their predicted functional impact.

**Table S5**. Differentially spliced events detected between embryonic and fetal hearts.

**Table S6**. Selected AS events between embryonic and fetal hearts, along with their predicted functional impact.

**Table S7**. Cardiomyocyte-enriched genes (genes with log2FC > 0.5 when comparing cardiomyocytes with all other cell types in the fetal heart single-cell atlas)

**Table S8**. Proteomic validation of developmentally regulated splice isoforms.

**Table S9**. Splicing events with divergent patterns in iPSC-CMs and hearts.

**Table S10**. Selected iPSC-CM-specific AS events, along with their predicted functional impact.

**Table S11**. RBPs differentially expressed between iPSC-CMs and both prenatal and postnatal hearts.

**Table S12**. AS-specific primers for quantitative PCR.

## References

1. Buijtendijk, M.F.J., Barnett, P., and Van Den Hoff, M.J.B. (2020). Development of the human heart. American J of Med Genetics Pt C 184, 7–22. 10.1002/ajmg.c.31778.

2. Luna-Zurita, L., Stirnimann, C.U., Glatt, S., Kaynak, B.L., Thomas, S., Baudin, F., Samee, M.A.H., He, D., Small, E.M., Mileikovsky, M., et al. (2016). Complex Interdependence Regulates Heterotypic Transcription Factor Distribution and Coordinates Cardiogenesis. Cell 164, 999–1014. 10.1016/j.cell.2016.01.004.

3. Olson, E.N. (2006). Gene Regulatory Networks in the Evolution and Development of the Heart. Science 313, 1922–1927. 10.1126/science.1132292.

4. Sylva, M., Van Den Hoff, M.J.B., and Moorman, A.F.M. (2014). Development of the human heart. American J of Med Genetics Pt A 164, 1347–1371. 10.1002/ajmg.a.35896.

5. Meilhac, S.M., and Buckingham, M.E. (2018). The deployment of cell lineages that form the mammalian heart. Nat Rev Cardiol 15, 705–724. 10.1038/s41569-018-0086-9.

6. Nilsen, T.W., and Graveley, B.R. (2010). Expansion of the eukaryotic proteome by alternative splicing. Nature 463, 457–463. 10.1038/nature08909.

7. Wang, E.T., Sandberg, R., Luo, S., Khrebtukova, I., Zhang, L., Mayr, C., Kingsmore, S.F., Schroth, G.P., and Burge, C.B. (2008). Alternative isoform regulation in human tissue transcriptomes. Nature 456, 470–476. 10.1038/nature07509.

8. Braunschweig, U., Barbosa-Morais, N.L., Pan, Q., Nachman, E.N., Alipanahi, B., Gonatopoulos-Pournatzis, T., Frey, B., Irimia, M., and Blencowe, B.J. (2014). Widespread intron retention in mammals functionally tunes transcriptomes. Genome Res. 24, 1774– 1786. 10.1101/gr.177790.114.

9. Kjer-Hansen, P., and Weatheritt, R.J. (2023). The function of alternative splicing in the proteome: rewiring protein interactomes to put old functions into new contexts. Nat Struct Mol Biol 30, 1844–1856. 10.1038/s41594-023-01155-9.

10. Yang, X., Coulombe-Huntington, J., Kang, S., Sheynkman, G.M., Hao, T., Richardson, A., Sun, S., Yang, F., Shen, Y.A., Murray, R.R., et al. (2016). Widespread Expansion of Protein Interaction Capabilities by Alternative Splicing. Cell 164, 805–817. 10.1016/j.cell.2016.01.029.

11. Baralle, F.E., and Giudice, J. (2017). Alternative splicing as a regulator of development and tissue identity. Nat Rev Mol Cell Biol 18, 437–451. 10.1038/nrm.2017.27.

12. Blencowe, B.J. (2006). Alternative Splicing: New Insights from Global Analyses. Cell 126, 37–47. 10.1016/j.cell.2006.06.023.

13. Merkin, J., Russell, C., Chen, P., and Burge, C.B. (2012). Evolutionary Dynamics of Gene and Isoform Regulation in Mammalian Tissues. Science 338, 1593–1599. 10.1126/science.1228186.

14. Jimena Giudice, Giudice, J., Zheng Xia, Xia, Z., Eric T. Wang, Wang, E.T., Eric T. Wang, Marissa A. Scavuzzo, Scavuzzo, M.A., Amanda J. Ward, et al. (2014). Alternative splicing regulates vesicular trafficking genes in cardiomyocytes during postnatal heart development. Nature Communications 5, 3603–3603. 10.1038/ncomms4603.

15. Kalsotra, A., Xiao, X., Ward, A.J., Castle, J.C., Johnson, J.M., Burge, C.B., and Cooper, T.A. (2008). A postnatal switch of CELF and MBNL proteins reprograms alternative splicing in the developing heart. Proc. Natl. Acad. Sci. U.S.A. 105, 20333–20338. 10.1073/pnas.0809045105.

16. Van Den Hoogenhof, M.M.G., Pinto, Y.M., and Creemers, E.E. (2016). RNA Splicing: Regulation and Dysregulation in the Heart. Circ Res 118, 454–468. 10.1161/CIRCRESAHA.115.307872.

17. Mazin, P.V., Khaitovich, P., Cardoso-Moreira, M., and Kaessmann, H. (2021). Alternative splicing during mammalian organ development. Nat Genet 53, 925–934. 10.1038/s41588-021-00851-w.

18. Burridge, P.W., Sharma, A., and Wu, J.C. (2015). Genetic and Epigenetic Regulation of Human Cardiac Reprogramming and Differentiation in Regenerative Medicine. Annu. Rev. Genet. 49, 461–484. 10.1146/annurev-genet-112414-054911.

19. Zaragoza, C., Gomez-Guerrero, C., Martin-Ventura, J.L., Blanco-Colio, L., Lavin, B., Mallavia, B., Tarin, C., Mas, S., Ortiz, A., and Egido, J. (2011). Animal Models of Cardiovascular Diseases. Journal of Biomedicine and Biotechnology 2011, 1–13. 10.1155/2011/497841.

20. Takahashi, K., Tanabe, K., Ohnuki, M., Narita, M., Ichisaka, T., Tomoda, K., and Yamanaka, S. (2007). Induction of Pluripotent Stem Cells from Adult Human Fibroblasts by Defined Factors. Cell 131, 861–872. 10.1016/j.cell.2007.11.019.

21. Karbassi, E., Fenix, A., Marchiano, S., Muraoka, N., Nakamura, K., Yang, X., and Murry, C.E. (2020). Cardiomyocyte maturation: advances in knowledge and implications for regenerative medicine. Nat Rev Cardiol 17, 341–359. 10.1038/s41569-019-0331-x.

22. Wang, J., An, M., Haubner, B.J., and Penninger, J.M. (2023). Cardiac regeneration: Options for repairing the injured heart. Front. Cardiovasc. Med. 9, 981982. 10.3389/fcvm.2022.981982.

23. Farah, E.N., Hu, R.K., Kern, C., Zhang, Q., Lu, T.-Y., Ma, Q., Tran, S., Zhang, B., Carlin, D., Monell, A., et al. (2024). Spatially organized cellular communities form the developing human heart. Nature. 10.1038/s41586-024-07171-z.

24. Cao, J., O’Day, D.R., Pliner, H.A., Kingsley, P.D., Deng, M., Daza, R.M., Zager, M.A., Aldinger, K.A., Blecher-Gonen, R., Zhang, F., et al. (2020). A human cell atlas of fetal gene expression. Science 370, eaba7721. 10.1126/science.aba7721.

25. Cui, Y., Zheng, Y., Liu, X., Yan, L., Fan, X., Yong, J., Hu, Y., Dong, J., Li, Q., Wu, X., et al. (2019). Single-Cell Transcriptome Analysis Maps the Developmental Track of the Human Heart. Cell Reports 26, 1934–1950.e5. 10.1016/j.celrep.2019.01.079.

26. Galdos, F.X., Lee, C., Lee, S., Paige, S., Goodyer, W., Xu, S., Samad, T., Escobar, G.V., Darsha, A., Beck, A., et al. (2023). Combined lineage tracing and scRNA-seq reveals unexpected first heart field predominance of human iPSC differentiation. eLife 12, e80075. 10.7554/eLife.80075.

27. Tyser, R.C.V., Ibarra-Soria, X., McDole, K., Arcot Jayaram, S., Godwin, J., Van Den Brand, T.A.H., Miranda, A.M.A., Scialdone, A., Keller, P.J., Marioni, J.C., et al. (2021). Characterization of a common progenitor pool of the epicardium and myocardium. Science 371, eabb2986. 10.1126/science.abb2986.

28. Cardoso-Moreira, M., Halbert, J., Valloton, D., Velten, B., Chen, C., Shao, Y., Liechti, A., Ascenção, K., Rummel, C., Ovchinnikova, S., et al. (2019). Gene expression across mammalian organ development. Nature 571, 505–509. 10.1038/s41586-019-1338-5.

29. Maroli, G., and Braun, T. (2021). The long and winding road of cardiomyocyte maturation. Cardiovascular Research 117, 712–726. 10.1093/cvr/cvaa159.

30. Vučković, S., Dinani, R., Nollet, E.E., Kuster, D.W.D., Buikema, J.W., Houtkooper, R.H., Nabben, M., Van Der Velden, J., and Goversen, B. (2022). Characterization of cardiac metabolism in iPSC-derived cardiomyocytes: lessons from maturation and disease modeling. Stem Cell Res Ther 13, 332. 10.1186/s13287-022-03021-9.

31. Jiang, X., Lian, X., Wei, K., Zhang, J., Yu, K., Li, H., Ma, H., Cai, Y., and Pang, L. (2024). Maturation of pluripotent stem cell-derived cardiomyocytes: limitations and challenges from metabolic aspects. Stem Cell Res Ther 15. 10.1186/s13287-024-03961-4.

32. Efthymiou, G., Radwanska, A., Grapa, A.-I., Beghelli-de La Forest Divonne, S., Grall, D., Schaub, S., Hattab, M., Pisano, S., Poet, M., Pisani, D.F., et al. (2021). Fibronectin Extra Domains tune cellular responses and confer topographically distinct features to fibril networks. Journal of Cell Science 134, jcs252957. 10.1242/jcs.252957.

33. Godbout, E., Son, D.O., Hume, S., Boo, S., Sarrazy, V., Clément, S., Kapus, A., Wehrle-Haller, B., Bruckner-Tuderman, L., Has, C., et al. (2020). Kindlin-2 Mediates Mechanical Activation of Cardiac Myofibroblasts. Cells 9, 2702. 10.3390/cells9122702.

34. Timpl, R., Sasaki, T., Kostka, G., and Chu, M.-L. (2003). Fibulins: a versatile family of extracellular matrix proteins. Nat Rev Mol Cell Biol 4, 479–489. 10.1038/nrm1130.

35. Gönczi, M., Teixeira, J.M.C., Barrera-Vilarmau, S., Mediani, L., Antoniani, F., Nagy, T.M., Fehér, K., Ráduly, Z., Ambrus, V., Tőzsér, J., et al. (2023). Alternatively spliced exon regulates context-dependent MEF2D higher-order assembly during myogenesis. Nat Commun 14, 1329. 10.1038/s41467-023-37017-7.

36. Yamazaki, T., Liu, L., Lazarev, D., Al-Zain, A., Fomin, V., Yeung, P.L., Chambers, S.M., Lu, C.-W., Studer, L., and Manley, J.L. (2018). TCF3 alternative splicing controlled by hnRNP H/F regulates E-cadherin expression and hESC pluripotency. Genes Dev. 32, 1161– 1174. 10.1101/gad.316984.118.

37. Yoneda-Kato, N., Tomoda, K., Umehara, M., Arata, Y., and Kato, J. (2005). Myeloid leukemia factor 1 regulates p53 by suppressing COP1 via COP9 signalosome subunit 3. EMBO J 24, 1739–1749. 10.1038/sj.emboj.7600656.

38. John, G.B., Shang, Y., Li, L., Renken, C., Mannella, C.A., Selker, J.M.L., Rangell, L., Bennett, M.J., and Zha, J. (2005). The Mitochondrial Inner Membrane Protein Mitofilin Controls Cristae Morphology. MBoC 16, 1543–1554. 10.1091/mbc.e04-08-0697.

39. Bock-Bierbaum, T., Funck, K., Wollweber, F., Lisicki, E., Von Der Malsburg, K., Von Der Malsburg, A., Laborenz, J., Noel, J.K., Hessenberger, M., Jungbluth, S., et al. (2022). Structural insights into crista junction formation by the Mic60-Mic19 complex. Sci. Adv. 8, eabo4946. 10.1126/sciadv.abo4946.

40. Christofk, H.R., Vander Heiden, M.G., Harris, M.H., Ramanathan, A., Gerszten, R.E., Wei, R., Fleming, M.D., Schreiber, S.L., and Cantley, L.C. (2008). The M2 splice isoform of pyruvate kinase is important for cancer metabolism and tumour growth. Nature 452, 230–233. 10.1038/nature06734.

41. Di Minin, G., Holzner, M., Grison, A., Dumeau, C.E., Chan, W., Monfort, A., Jerome-Majewska, L.A., Roelink, H., and Wutz, A. (2022). TMED2 binding restricts SMO to the ER and Golgi compartments. PLoS Biol 20, e3001596. 10.1371/journal.pbio.3001596.

42. Gomes, A.V., Guzman, G., Zhao, J., and Potter, J.D. (2002). Cardiac troponin T isoforms affect the Ca2+ sensitivity and inhibition of force development. Insights into the role of troponin T isoforms in the heart. J Biol Chem 277, 35341–35349. 10.1074/jbc.M204118200.

43. Lu, F., Ma, Q., Xie, W., Liou, C.L., Zhang, D., Sweat, M.E., Jardin, B.D., Naya, F.J., Guo, Y., Cheng, H., et al. (2022). CMYA5 establishes cardiac dyad architecture and positioning. Nat Commun 13, 2185. 10.1038/s41467-022-29902-4.

44. Huang, C., Zhou, Q., Liang, P., Hollander, M.S., Sheikh, F., Li, X., Greaser, M., Shelton, G.D., Evans, S., and Chen, J. (2003). Characterization and in Vivo Functional Analysis of Splice Variants of Cypher. Journal of Biological Chemistry 278, 7360–7365. 10.1074/jbc.M211875200.

45. Chopra, N., and Knollmann, B.C. (2013). Triadin regulates cardiac muscle couplon structure and microdomain Ca2+ signalling: a path towards ventricular arrhythmias. Cardiovascular Research 98, 187–191. 10.1093/cvr/cvt023.

46. Duran, J., Nickel, L., Estrada, M., Backs, J., and Van Den Hoogenhof, M.M.G. (2021). CaMKIIδ Splice Variants in the Healthy and Diseased Heart. Front. Cell Dev. Biol. 9, 644630. 10.3389/fcell.2021.644630.

47. Doll, S., Dreßen, M., Geyer, P.E., Itzhak, D.N., Braun, C., Doppler, S.A., Meier, F., Deutsch, M.-A., Lahm, H., Lange, R., et al. (2017). Region and cell-type resolved quantitative proteomic map of the human heart. Nat Commun 8, 1469. 10.1038/s41467-017-01747-2.

48. Colwill, K., Pawson, T., Andrews, B., Prasad, J., Manley, J.L., Bell, J.C., and Duncan, P.I. (1996). The Clk/Sty protein kinase phosphorylates SR splicing factors and regulates their intranuclear distribution. EMBO J 15, 265–275.

49. Guo, Y., and Pu, W.T. (2020). Cardiomyocyte Maturation: New Phase in Development. Circulation Research 126, 1086–1106. 10.1161/CIRCRESAHA.119.315862.

50. Synnergren, J., Améen, C., Jansson, A., and Sartipy, P. (2012). Global transcriptional profiling reveals similarities and differences between human stem cell-derived cardiomyocyte clusters and heart tissue. Physiological Genomics 44, 245–258. 10.1152/physiolgenomics.00118.2011.

51. Van Den Berg, C.W., Okawa, S., Chuva De Sousa Lopes, S.M., Van Iperen, L., Passier, R., Braam, S.R., Tertoolen, L.G., Del Sol, A., Davis, R.P., and Mummery, C.L. (2015). Transcriptome of human foetal heart compared with cardiomyocytes from pluripotent stem cells. Development, dev.123810. 10.1242/dev.123810.

52. Giudice, J., and Cooper, T.A. (2014). RNA-Binding Proteins in Heart Development. In Systems Biology of RNA Binding Proteins Advances in Experimental Medicine and Biology., G. W. Yeo, ed. (Springer New York), pp. 389–429. 10.1007/978-1-4939-1221-6_11.

53. Li, Z., Cao, C., Zhao, Q., Li, D., Han, Y., Zhang, M., Mao, L., Zhou, B., and Wang, L. (2024). RNA splicing controls organ-wide maturation of postnatal heart in mice. Developmental Cell 0. 10.1016/j.devcel.2024.09.018.

54. Weeland, C.J., Van Den Hoogenhof, M.M., Beqqali, A., and Creemers, E.E. (2015). Insights into alternative splicing of sarcomeric genes in the heart. Journal of Molecular and Cellular Cardiology 81, 107–113. 10.1016/j.yjmcc.2015.02.008.

55. Agarkova, I., Auerbach, D., Ehler, E., and Perriard, J.C. (2000). A novel marker for vertebrate embryonic heart, the EH-myomesin isoform. J Biol Chem 275, 10256–10264. 10.1074/jbc.275.14.10256.

56. Wang, E.T., Cody, N.A.L., Jog, S., Biancolella, M., Wang, T.T., Treacy, D.J., Luo, S., Schroth, G.P., Housman, D.E., Reddy, S., et al. (2012). Transcriptome-wide Regulation of Pre-mRNA Splicing and mRNA Localization by Muscleblind Proteins. Cell 150, 710–724. 10.1016/j.cell.2012.06.041.

57. Xu, X., Yang, D., Ding, J.-H., Wang, W., Chu, P.-H., Dalton, N.D., Wang, H.-Y., Bermingham, J.R., Ye, Z., Liu, F., et al. (2005). ASF/SF2-regulated CaMKIIdelta alternative splicing temporally reprograms excitation-contraction coupling in cardiac muscle. Cell 120, 59–72. 10.1016/j.cell.2004.11.036.

58. Cao, J., Routh, A.L., and Kuyumcu-Martinez, M.N. (2021). Nanopore sequencing reveals full-length Tropomyosin 1 isoforms and their regulation by RNA-binding proteins during rat heart development. J Cell Mol Med 25, 8352–8362. 10.1111/jcmm.16795.

59. Charlet-B, N., Singh, G., Cooper, T.A., and Logan, P. (2002). Dynamic Antagonism between ETR-3 and PTB Regulates Cell Type-Specific Alternative Splicing. Molecular Cell 9, 649– 658. 10.1016/S1097-2765(02)00479-3.

60. Verma, S.K., Deshmukh, V., Thatcher, K., Belanger, K.K., Rhyner, A.M., Meng, S., Holcomb, R.J., Bressan, M., Martin, J.F., Cooke, J.P., et al. (2022). RBFOX2 is required for establishing RNA regulatory networks essential for heart development. Nucleic Acids Res 50, 2270–2286. 10.1093/nar/gkac055.

61. Zhang, X., Wang, Z., Xu, Q., Chen, Y., Liu, W., Zhong, T., Li, H., Quan, C., Zhang, L., and Cui, C.-P. (2021). Splicing factor Srsf5 deletion disrupts alternative splicing and causes noncompaction of ventricular myocardium. iScience 24, 103097. 10.1016/j.isci.2021.103097.

62. Feng, Y., Valley, M.T., Lazar, J., Yang, A.L., Bronson, R.T., Firestein, S., Coetzee, W.A., and Manley, J.L. (2009). SRp38 regulates alternative splicing and is required for Ca(2+) handling in the embryonic heart. Dev Cell 16, 528–538. 10.1016/j.devcel.2009.02.009.

63. Poon, K.L., Tan, K.T., Wei, Y.Y., Ng, C.P., Colman, A., Korzh, V., and Xu, X.Q. (2012). RNA-binding protein RBM24 is required for sarcomere assembly and heart contractility. Cardiovasc Res 94, 418–427. 10.1093/cvr/cvs095.

64. Yang, J., Hung, L.-H., Licht, T., Kostin, S., Looso, M., Khrameeva, E., Bindereif, A., Schneider, A., and Braun, T. (2014). RBM24 is a major regulator of muscle-specific alternative splicing. Dev Cell 31, 87–99. 10.1016/j.devcel.2014.08.025.

65. Guo, W., Schafer, S., Greaser, M.L., Radke, M.H., Liss, M., Govindarajan, T., Maatz, H., Schulz, H., Li, S., Parrish, A.M., et al. (2012). RBM20, a gene for hereditary cardiomyopathy, regulates titin splicing. Nat Med 18, 766–773. 10.1038/nm.2693.

66. Li, S., Guo, W., Dewey, C.N., and Greaser, M.L. (2013). Rbm20 regulates titin alternative splicing as a splicing repressor. Nucleic Acids Res 41, 2659–2672. 10.1093/nar/gks1362.

67. Montañés-Agudo, P., Aufiero, S., Schepers, E.N., van der Made, I., Pinto, Y.M., and Creemers, E. (2022). The RNA-binding protein QKI governs a muscle-specific alternative splicing program that shapes the contractile function of cardiomyocytes. Journal of Molecular and Cellular Cardiology 173, S161. 10.1016/j.yjmcc.2022.08.317.

68. Zhong, X.-Y., Ding, J.-H., Adams, J.A., Ghosh, G., and Fu, X.-D. (2009). Regulation of SR protein phosphorylation and alternative splicing by modulating kinetic interactions of SRPK1 with molecular chaperones. Genes Dev. 23, 482–495. 10.1101/gad.1752109.

69. Töpf, A., Cox, D., Zaharieva, I.T., Di Leo, V., Sarparanta, J., Jonson, P.H., Sealy, I.M., Smolnikov, A., White, R.J., Vihola, A., et al. (2024). Digenic inheritance involving a muscle-specific protein kinase and the giant titin protein causes a skeletal muscle myopathy. Nat Genet 56, 395–407. 10.1038/s41588-023-01651-0.

70. Murayama, R., Kimura-Asami, M., Togo-Ohno, M., Yamasaki-Kato, Y., Naruse, T.K., Yamamoto, T., Hayashi, T., Ai, T., Spoonamore, K.G., Kovacs, R.J., et al. (2018). Phosphorylation of the RSRSP stretch is critical for splicing regulation by RNA-Binding Motif Protein 20 (RBM20) through nuclear localization. Sci Rep 8, 8970. 10.1038/s41598-018-26624-w.

71. Sun, M., Jin, Y., Zhang, Y., Gregorich, Z.R., Ren, J., Ge, Y., and Guo, W. (2022). SR Protein Kinases Regulate the Splicing of Cardiomyopathy-Relevant Genes via Phosphorylation of the RSRSP Stretch in RBM20. Genes 13, 1526. 10.3390/genes13091526.

72. Yu, J., Hu, K., Smuga-Otto, K., Tian, S., Stewart, R., Slukvin, I.I., and Thomson, J.A. (2009). Human Induced Pluripotent Stem Cells Free of Vector and Transgene Sequences. Science 324, 797–801. 10.1126/science.1172482.

73. Burridge, P.W., Matsa, E., Shukla, P., Lin, Z.C., Churko, J.M., Ebert, A.D., Lan, F., Diecke, S., Huber, B., Mordwinkin, N.M., et al. (2014). Chemically defined generation of human cardiomyocytes. Nat Methods 11, 855–860. 10.1038/nmeth.2999.

74. Knight, W.E., Cao, Y., Lin, Y.-H., Chi, C., Bai, B., Sparagna, G.C., Zhao, Y., Du, Y., Londono, P., Reisz, J.A., et al. (2021). Maturation of Pluripotent Stem Cell-Derived Cardiomyocytes Enables Modeling of Human Hypertrophic Cardiomyopathy. Stem Cell Reports 16, 519–533. 10.1016/j.stemcr.2021.01.018.

75. Silva, T.P., Bekman, E.P., Fernandes, T.G., Vaz, S.H., Rodrigues, C.A.V., Diogo, M.M., Cabral, J.M.S., and Carmo-Fonseca, M. (2020). Maturation of Human Pluripotent Stem Cell-Derived Cerebellar Neurons in the Absence of Co-culture. Front. Bioeng. Biotechnol. 8, 70. 10.3389/fbioe.2020.00070.

76. Moretti, A., Bellin, M., Welling, A., Jung, C.B., Lam, J.T., Bott-Flügel, L., Dorn, T., Goedel, A., Höhnke, C., Hofmann, F., et al. (2010). Patient-Specific Induced Pluripotent Stem-Cell Models for Long-QT Syndrome. N Engl J Med 363, 1397–1409. 10.1056/NEJMoa0908679.

77. Jager, J., Ribeiro, M., Furtado, M., Carvalho, T., Syrris, P., Lopes, L.R., Elliott, P.M., Cabral, J.M.S., Carmo-Fonseca, M., Da Rocha, S.T., et al. (2024). Patient-derived induced pluripotent stem cells to study non-canonical splicing variants associated with Hypertrophic Cardiomyopathy. Stem Cell Research 81, 103582. 10.1016/j.scr.2024.103582.

78. Breckwoldt, K., Letuffe-Brenière, D., Mannhardt, I., Schulze, T., Ulmer, B., Werner, T., Benzin, A., Klampe, B., Reinsch, M.C., Laufer, S., et al. (2017). Differentiation of cardiomyocytes and generation of human engineered heart tissue. Nat Protoc 12, 1177– 1197. 10.1038/nprot.2017.033.

79. Radke, M.H., Badillo-Lisakowski, V., Britto-Borges, T., Kubli, D.A., Jüttner, R., Parakkat, P., Carballo, J.L., Hüttemeister, J., Liss, M., Hansen, A., et al. (2021). Therapeutic inhibition of RBM20 improves diastolic function in a murine heart failure model and human engineered heart tissue. Sci. Transl. Med. 13, eabe8952. 10.1126/scitranslmed.abe8952.

80. Melsted, P., Booeshaghi, A.S., Liu, L., Gao, F., Lu, L., Min, K.H., Da Veiga Beltrame, E., Hjörleifsson, K.E., Gehring, J., and Pachter, L. (2021). Modular, efficient and constant-memory single-cell RNA-seq preprocessing. Nat Biotechnol 39, 813–818. 10.1038/s41587-021-00870-2.

81. Hao, Y., Hao, S., Andersen-Nissen, E., Mauck, W.M., Zheng, S., Butler, A., Lee, M.J., Wilk, A.J., Darby, C., Zager, M., et al. (2021). Integrated analysis of multimodal single-cell data. Cell 184, 3573–3587.e29. 10.1016/j.cell.2021.04.048.

82. participants in the 1st Human Cell Atlas Jamboree, Lun, A.T.L., Riesenfeld, S., Andrews, T., Dao, T.P., Gomes, T., and Marioni, J.C. (2019). EmptyDrops: distinguishing cells from empty droplets in droplet-based single-cell RNA sequencing data. Genome Biol 20, 63. 10.1186/s13059-019-1662-y.

83. Mercer, T.R., Neph, S., Dinger, M.E., Crawford, J., Smith, M.A., Shearwood, A.-M.J., Haugen, E., Bracken, C.P., Rackham, O., Stamatoyannopoulos, J.A., et al. (2011). The Human Mitochondrial Transcriptome. Cell 146, 645–658. 10.1016/j.cell.2011.06.051.

84. Wu, T., Hu, E., Xu, S., Chen, M., Guo, P., Dai, Z., Feng, T., Zhou, L., Tang, W., Zhan, L., et al. (2021). clusterProfiler 4.0: A universal enrichment tool for interpreting omics data. The Innovation 2, 100141. 10.1016/j.xinn.2021.100141.

85. Carlson, M. (2017). org.Hs.eg.db. (Bioconductor). 10.18129/B9.BIOC.ORG.HS.EG.DB https://doi.org/10.18129/B9.BIOC.ORG.HS.EG.DB.

86. Andrews S (2010). FastQC: A Quality Control Tool for High Throughput Sequence Data.

87. Krueger, F., James, F., Ewels, P., Afyounian, E., and Schuster-Boeckler, B. (2021). FelixKrueger/TrimGalore: v0.6.7 - DOI via Zenodo. Version 0.6.7 (Zenodo). 10.5281/ZENODO.5127899 https://doi.org/10.5281/ZENODO.5127899.

88. Patro, R., Duggal, G., Love, M.I., Irizarry, R.A., and Kingsford, C. (2017). Salmon provides fast and bias-aware quantification of transcript expression. Nat Methods 14, 417–419. 10.1038/nmeth.4197.

89. Dobin, A., Davis, C.A., Schlesinger, F., Drenkow, J., Zaleski, C., Jha, S., Batut, P., Chaisson, M., and Gingeras, T.R. (2013). STAR: ultrafast universal RNA-seq aligner. Bioinformatics 29, 15–21. 10.1093/bioinformatics/bts635.

90. Li, H., Handsaker, B., Wysoker, A., Fennell, T., Ruan, J., Homer, N., Marth, G., Abecasis, G., Durbin, R., and 1000 Genome Project Data Processing Subgroup (2009). The Sequence Alignment/Map format and SAMtools. Bioinformatics 25, 2078–2079. 10.1093/bioinformatics/btp352.

91. Frankish, A., Diekhans, M., Ferreira, A.-M., Johnson, R., Jungreis, I., Loveland, J., Mudge, J.M., Sisu, C., Wright, J., Armstrong, J., et al. (2019). GENCODE reference annotation for the human and mouse genomes. Nucleic Acids Research 47, D766–D773. 10.1093/nar/gky955.

92. R Core Team (2021). R: A language and environment for statistical computing. (R Foundation for Statistical Computing, Vienna, Austria.).

93. Soneson, C., Love, M.I., and Robinson, M.D. (2016). Differential analyses for RNA-seq: transcript-level estimates improve gene-level inferences. F1000Res 4, 1521. 10.12688/f1000research.7563.2.

94. Wei, Taiyun and Simko, Viktor R package “corrplot”: Visualization of a Correlation Matrix.

95. Chen, H., and Boutros, P.C. (2011). VennDiagram: a package for the generation of highly-customizable Venn and Euler diagrams in R. BMC Bioinformatics 12, 35. 10.1186/1471-2105-12-35.

96. Love, M.I., Huber, W., and Anders, S. (2014). Moderated estimation of fold change and dispersion for RNA-seq data with DESeq2. Genome Biol 15, 550. 10.1186/s13059-014-0550-8.

97. Wickham, H. (2016). ggplot2 (Springer International Publishing) 10.1007/978-3-319-24277-4.

98. Suzuki, R., and Shimodaira, H. (2006). Pvclust: an R package for assessing the uncertainty in hierarchical clustering. Bioinformatics 22, 1540–1542. 10.1093/bioinformatics/btl117.

99. Igor Dolgalev (2025). msigdbr: MSigDB Gene Sets for Multiple Organisms in a Tidy Data Format. Version 25.1.0.

100. Hänzelmann, S., Castelo, R., and Guinney, J. (2013). GSVA: gene set variation analysis for microarray and RNA-Seq data. BMC Bioinformatics 14. 10.1186/1471-2105-14-7.

101. Newman, A.M., Steen, C.B., Liu, C.L., Gentles, A.J., Chaudhuri, A.A., Scherer, F., Khodadoust, M.S., Esfahani, M.S., Luca, B.A., Steiner, D., et al. (2019). Determining cell type abundance and expression from bulk tissues with digital cytometry. Nat Biotechnol 37, 773–782. 10.1038/s41587-019-0114-2.

102. Shen, S., Park, J.W., Lu, Z., Lin, L., Henry, M.D., Wu, Y.N., Zhou, Q., and Xing, Y. (2014). rMATS: Robust and flexible detection of differential alternative splicing from replicate RNA-Seq data. Proc. Natl. Acad. Sci. U.S.A. 111. 10.1073/pnas.1419161111.

103. Vaquero-Garcia, J., Aicher, J.K., Jewell, S., Gazzara, M.R., Radens, C.M., Jha, A., Norton, S.S., Lahens, N.F., Grant, G.R., and Barash, Y. (2023). RNA splicing analysis using heterogeneous and large RNA-seq datasets. Nat Commun 14, 1230. 10.1038/s41467-023-36585-y.

104. Tapial, J., Ha, K.C.H., Sterne-Weiler, T., Gohr, A., Braunschweig, U., Hermoso-Pulido, A., Quesnel-Vallières, M., Permanyer, J., Sodaei, R., Marquez, Y., et al. (2017). An atlas of alternative splicing profiles and functional associations reveals new regulatory programs and genes that simultaneously express multiple major isoforms. Genome Res. 27, 1759– 1768. 10.1101/gr.220962.117.

105. Kolde R (2019). pheatmap: Pretty Heatmaps.

106. Neuwirth E (2014). RColorBrewer: ColorBrewer Palettes.

107. Garrido-Martín, D., Palumbo, E., Guigó, R., and Breschi, A. (2018). ggsashimi: Sashimi plot revised for browser- and annotation-independent splicing visualization. PLoS Comput Biol 14, e1006360. 10.1371/journal.pcbi.1006360.

108. Louadi, Z., Elkjaer, M.L., Klug, M., Lio, C.T., Fenn, A., Illes, Z., Bongiovanni, D., Baumbach, J., Kacprowski, T., List, M., et al. (2021). Functional enrichment of alternative splicing events with NEASE reveals insights into tissue identity and diseases. Genome Biol 22, 327. 10.1186/s13059-021-02538-1.

109. Maia, T.M., Staes, A., Plasman, K., Pauwels, J., Boucher, K., Argentini, A., Martens, L., Montoye, T., Gevaert, K., and Impens, F. (2020). Simple Peptide Quantification Approach for MS-Based Proteomics Quality Control. ACS Omega 5, 6754–6762. 10.1021/acsomega.0c00080.

110. Chiva, C., Olivella, R., Borràs, E., Espadas, G., Pastor, O., Solé, A., and Sabidó, E. (2018). QCloud: A cloud-based quality control system for mass spectrometry-based proteomics laboratories. PLoS ONE 13, e0189209. 10.1371/journal.pone.0189209.

