## Supplementary material for "Alternative splicing dynamics during human cardiac development *in vivo* and *in vitro*": Table S4

| **Functional category** | **Gene** | **Exon** | **PSI Prenatal Hearts** | **PSI Postnatal Hearts** | **Affected domain/ motif** |
| --- | --- | --- | --- | --- | --- |
| Metabolism | ***ACSS2*** | chr20:34915218-34915256 | 0.43 | 0.87 | AMP-binding domain (PF00501) |
| Muscle Contraction | ***ACTN4*** | chr19:38710257-38710342 | 0.59 | 0.83 | Calponin homology (CH) domain (PF00307) |
| Ion Channel | ***ANK2*** | chr4:113378090-113378181 | 0.16 | 0.54 | Disordered region |
| Ion Channel | ***CAMK2D*** | chr4:113455726-113455821 | 0.46 | 0.88 | Disordered region |
| Sarcomeric | ***CMYA5*** | chr5:79747091-79747113 | 0.21 | 0.99 | ORF disruption upon exon exclusion  GLU_RICH  Disordered region |
| Cytoskeleton | ***COBL*** | chr7:51187914-51187958 | 0.02 | 0.32 | Cordon-bleu ubiquitin-like domain (PF09469)  Disordered region |
| ECM organization | ***DCN*** | chr12:91179362-91179686 | 0.03 | 0.31 | - |
| Transcription | ***DMPK*** | chr19:45771350-45771396 | 0.95 | 0.74 | - |
| ECM organization | ***FBLN2*** | chr3:13621775-13621915 | 0.08 | 0.46 | Calcium-binding EGF domain (PF00307) |
| ECM/ Cell-cell communication | ***FERMT2*** | chr14:52861587-52861610 | 0.04 | 0.43 | FERM central domain (PF00373) |
| ECM | ***FGFR1*** | chr8:38429682-38429948 | 0.68 | 0.38 | Immunoglobulin (Ig) domain (PF00047) |
| ECM | ***FN1*** | chr2:215392931-215393203 | 0.46 | 0.11 | Fibronectin type III domain (PF00041)  Disordered region |
| ECM | ***FN1*** | chr2:215380811-215381080 | 0.48 | 0.11 | Fibronectin type III domain (PF00041)  Disordered region |
| Translation | ***FXR1*** | chr3:180971075-180971155 | 0.55 | 0.80 | - |
| Transcription | ***HMGN3*** | chr6:79202063-79202155 | 0.31 | 0.68 | Nucleosomal binding domain (PF01101)  Disordered region |
| Mitochondrial | ***IDH3B*** | chr20:2658395-2658522 | 0.40 | 0.18 | Isocitrate/isopropylmalate dehydrogenase domain (PF00180) |
| Mitochondrial | ***IMMT*** | chr2:86170749-86170841 | 0.57 | 0.18 | PF097314 Mitofilin  Disordered region |
| Transcription | ***IWS1*** | chr2:127485874-127486033 | 0.13 | 0.52 | Disordered region |
| Sarcomeric | ***LDB3*** | chr10:86706531-86706719 | 0.70 | 0.23 | Disordered region PS50099 PRO_RICH PS50310 ALA_RICH |
| Sarcomeric | ***LDB3*** | chr10:86681436-86681803 | 0.98 | 0.51 | - |
| Sarcomeric | ***LDB3*** | chr10:86685678-86685700 | 0 | 0.38 | Disordered region |
| Sarcomeric | ***LDB3*** | chr10:86687069-86687272 | 0.09 | 0.55 | Domain of unknown function (PF15936) Often contains Zasp-like motif |
| Histone | ***MACROH2A1*** | chr5:135352946-135353045 | 0.42 | 0.15 | ORF disruption upon sequence exclusion  Macro domain  (PF0166116). Binds ADP-ribose |
| Splicing | ***MBNL1*** | chr3:152446704-152446757 | 0.47 | 0.12 | PS50310 Ala-Rich |
| Transcription | ***MEF2D*** | chr1:156476494-156476514 | 0.09 | 0.46 | Disordered region |
| Mitochondrial | ***MFF*** | chr2:227342745-227342819 | 0.37 | 0.58 | Miff domain (PF05644) Disordered region |
| Cell cycle | ***MLF1*** | chr3:158593382-158593426 | 0.06 | 0.49 | Mlf1IP domain (PF10248)  Disordered region |
| Sarcomeric | ***MYL6*** | chr12:56160626-56160670 | 0.36 | 0.82 | - |
| Sarcomeric | ***MYO1B*** | chr2:191400749-191400835 | 0.57 | 0.12 | PS50096 IQ |
| Sarcomeric | ***MYO1B*** | chr2:191402632-191402718 | 0.59 | 0.29 | PS50096 IQ |
| Sarcomeric | ***MYOM1*** | chr18:3129232-3129519 | 0.75 | 0.01 | Disordered region between FN3 domains |
| ECM | ***NCAM1*** | chr11:113242805-113242846 | 0.07 | 0.76 | - |
| Transcription | ***NCOR2*** | chr12:124327547-124327633 | 0.83 | 0.41 | - |
| Mitochondrial | ***PKM*** | chr15:72203022-72203188 | 0.28 | 0.57 | Alpha/beta domain (PF02887), Barrel domain (PF00224) |
| Endosomal trafficking | ***PLEKHM2*** | chr1:15721329-15721388 | 0.72 | 0.98 | Binding motifs of TPR domain of kinesin light chain 1  Disordered region |
| Mitochondrial | ***SLC25A3*** | chr12:98595433-98595557 | 0.60 | 0.87 | Mitochondrial carrier protein  PF00153 |
| Muscle Contraction | ***SLMAP*** | chr3:57907884-57908006 | 0.38 | 0.14 | Filament domain (PF00038)  Disordered region |
| Mitochondrial | ***STAU2*** | chr8:73527691-73527804 | 0.29 | 0.63 | Staufen C-terminal domain (PF16482) |
| Transcription | ***TAF1*** | chrX:71459150-71459251 | 0.35 | 0.14 | - |
| Transcription | ***TCF3*** | chr19:1612207-1612430 | 0.34 | 0.63 | Helix-loop-helix DNA-binding domain (PF00010)  Co-resolved interactions with TAL1, LMO2, ID1 and NEUROD1 |
| Transcription | ***THOC5*** | chr22:29531078-29531110 | 0.38 | 0.81 | PF09766 |
| Vesicular Transport | ***TMED2*** | chr12:123587611-123587631 | 0.30 | 0.56 | GOLD-like domain (PF01105) |
| Muscle Contraction | ***TNNT2*** | chr1:201362386-201362391 | 0.25 | 0.89 | Troponin domain (PF00992) |
| Muscle Contraction | ***TNNT2*** | chr1:201369816-201369845 | 0.94 | 0.61 | - |
| Muscle Contraction | ***TPM3*** | chr1:154172029-154172104 | 0.71 | 0.37 | Coiled coil containing alternating actin binding sites (PF00261) |
| Ion Channel | ***TRDN*** | chr6:123375605-123375631 | 0.42 | 0.85 | LYS_RICH Disordered region |
| Vesicular Transport | ***USO1*** | chr4:75795336-75795356 | 0.04 | 0.49 | Uso1/p115 like vesicle tethering protein, head region (PF04869) |
| Metabolism | ***VDAC3*** | chr8:42396678-42396680 | 0.57 | 0.83 | Porin (PF01459) |
