## Supplementary material for "Alternative splicing dynamics during human cardiac development *in vivo* and *in vitro*": Table S6

| **Functional Category** | **Gene** | **Exon** | **PSI**  **1 month Heart** | **PSI**  **5 month Heart** | **Affected domain / motif** |
| --- | --- | --- | --- | --- | --- |
| Cytoskeleton | ***ABI1*** | chr10:26771075-26771089 | 0.36 | 0.79 | Abi-interactor HHR (PF07815) |
| Scaffolding | ***ANK2*** | chr4:113323759-113323794 | 0.23 | 0.55 | ZU5 domain (PF00791, PS51145) |
| Scaffolding | ***ANK3*** | chr10:60196145-60196243 | 0.42 | 0.12 | Ankyrin repeats (PF13637, PS50088) |
| Chromatin | ***ARID1B*** | chr6:157174847-157175005 | 0.74 | 0.96 | Serine rich region (PS50324)  Highly disordered region |
| Ion | ***ATP2B4*** | chr1:203702044-203702079 | 0.73 | 0.38 | E1-E2 ATPase (PF00122)  Highly disordered region |
| Transcription | ***DCAF6*** | chr1:168050892-168050933 | 0.08 | 0.29 | Nuclear localization signal (PS50079)  IQ motif (PS50096)  Highly disordered region |
| Cytoskeleton | ***EML1*** | chr14:99892139-99892195 | 0.08 | 0.58 | Hydrophobic EMAP-Like Protein (HELP) motif (PF03451) |
| Extracellular Matrix | ***FXR1*** | chr3:180971075-180971155 | 0.50 | 0.86 | Nuclear localization signal (PS50079)  Arginine rich region (PS50323)  Highly disordered region |
| Histone | ***H2AFY*** | chr5:135350823-135350913 | 0.47 | 0.79 | ORF disruption upon sequence inclusion  Macro domain (PF01661, PS51154) |
| Histone | ***H2AFY*** | chr5:135352946-135353045 | 0.57 | 0.24 | ORF disruption upon sequence exclusion  Macro domain (PF01661, PS51154) |
| Mitochondrial | ***IDH3B*** | chr20:2658638-2658837 | 0.40 | 0.11 | Isocitrate/isopropylmalate dehydrogenase domain (PF00180) |
| ECM | ***LAMA2*** | chr6:129292803-129293066 | 0.18 | 0.52 | Laminin EGF domain (PF00053, PS50027)  Cysteine-rich region (PS50311) |
| ECM | ***LAMA2*** | chr6:129443063-129443068 | 0.62 | 1 | Laminin Domain II (PF06009)  Highly disordered region |
| ECM | ***LAMA2*** | chr6:129475390-129475401 | 0.44 | 0.86 | Laminin G domain (PF00054, PS50025) |
| Cytoskeleton | ***MACF1*** | chr1:39465095-39465112 | 0.52 | 0.76 | Growth-Arrest-Specific Protein 2 (GAS2) Domain (GAS2) (PF02187, PS51460) |
| Splicing | ***MBNL2*** | chr13:97356796-97356849 | 0.60 | 0.17 | Highly disordered region |
| Microtubules | ***MDM1*** | chr12:68316581-68316610 | 0.23 | 0.66 | MDM1 domain (PF15501)  Highly disordered region |
| Cytoskeleton | ***MYO5A*** | chr15:52331699-52331836 | 0 | 0.37 | ORF disruption upon sequence inclusion  Highly disordered region |
| Cytoskeleton | ***PDLIM7*** | chr5:177491404-177491420 | 0.41 | 0.68 | ORF disruption upon sequence inclusion  Highly disordered region |
| mRNA processing | ***RBM26*** | chr13:79353153-79353224 | 0.24 | 0.72 | Highly disordered region |
| Cytoskeleton | ***SLAIN2*** | chr4:48394576-48394653 | 0.27 | 0.66 | SLAIN motif-containing family (PF15301)  Highly disordered region |
| Mitochondrial | ***SLC25A3*** | chr12:98595433-98595557 | 0.50 | 0.94 | Mitochondrial carrier protein  (PF00153, PS50920) |
| Mitochondrial | ***SLC25A3*** | chr12:98595727-98595848 | 0.48 | 0.07 | ORF disruption upon sequence exclusion  Mitochondrial carrier protein  (PF00153, PS50920) |
| Cell Cycle | ***SYNE1*** | chr6:152168113-152168281 | 0 | 0.23 | Spectrin repeat (PF00435) |
| Muscle Contraction | ***TPM3*** | chr1:154172029-154172104 | 0.92 | 0.59 | Tropomyosin (PF00261) |
| Ion Channel | ***TRDN*** | chr6:123269849-123269866 | 0.38 | 0.91 | Lysine-rich region (PS50318)  Highly disordered region |
| Angiogenesis | ***VEGFA*** | chr6:43780732-43780803 | 0.14 | 0.47 | VEGF heparin-binding domain (PF14554) |
