## Supplementary material for "Alternative splicing dynamics during human cardiac development *in vivo* and *in vitro*": Table S8

| **Gene Names** | **Samples** | **Splicing Events** | **Genome Coordinates** | **Peptides (MS/MS counts ratio)** | **PSI** |
| --- | --- | --- | --- | --- | --- |
| *MYOM1* | iPSC-CMs  (*n* = 3) | Exon 18 inclusion | chr18:3129232-3129519 | AAIGGGVSPDVCPALSDEPGGLTASR; VSETVQEELTPPPQK  *N* = 4 (82) | 0.95 |
|  |  | Exon 18 skipping |  | -  *N* = 0 (82) |  |
|  | Adult Heart  (*n* = 18) | Exon 18 inclusion |  | AAIGGGVSPDVCPALSDEPGGLTASR;  VHEASPPTFQK  *N* = 14 (12355) | 0.01 |
|  |  | Exon 18 skipping |  | AAIAPPSPPCDITCLESFR; AAIAPPSPPCDITCLESFRDSMVLGWK  *N* = 116 (12355) |  |
| *ACTN4* | iPSC-CMs  (*n* = 3) | Exon 8 inclusion | chr19:38710257- 38710342 | MLDAEDIVNTARPDEK  *N* = 2 (46) | 0.51 |
|  |  | Exon 8 skipping |  | -  *N* = 0 (46) |  |
|  | Adult Heart  (*n* = 18) | Exon 8 inclusion |  | MLDAEDIVNTARPDEK  *N* = 132 (2580) | 0.83 |
|  |  | Exon 8 skipping |  | -  *N* = 0 (2580) |  |
| *HMGN3* | iPSC-CMs  (*n* = 3) | Exon 6 inclusion | chr6:79202063-79202155 | STVNVSTSR  *N* = 2 (6) | 0.31 |
|  |  | Exon 6 skipping |  | AEEAQKTESVDNEGE  *N* = 1 (6) |  |
|  | Adult Heart  (*n* = 18) | Exon 6 inclusion |  | AEEIHISR; STVNVSTSR  *N* = 53 (53) | 0.68 |
|  |  | Exon 6 skipping |  | -  *N* = 0 (53) |  |
| *CAMK2D* | iPSC-CMs  (*n* = 3) | Exon 20 inclusion | chr4:113455726- 113455821 | SGSPTVPIK; ENFSGGTSLWQNI  *N* = 3 (24) | 0.34 |
|  |  | Exon 20 skipping |  | -  *N* = 0 (24) |  |
|  | Adult Heart  (*n* = 18) | Exon 20 inclusion |  | SGSPTVPIK  *N* = 20 (948) | 0.88 |
|  |  | Exon 20 skipping |  | SGSPTVPIN  *N* = 1 (948) |  |

**Table S7. Proteomic validation of developmentally regulated splice isoforms.**

For the indicated alternative splicing (AS) events, peptides uniquely corresponding to either the exon-inclusion or exon-skipping isoform are shown. *N* indicates the number of MS/MS spectra supporting each isoform-specific peptide; total spectral counts for the corresponding protein are shown in brackets. PSI represents the exon inclusion level derived from RNA-seq data.
