## Supplementary material for "Alternative splicing dynamics during human cardiac development *in vivo* and *in vitro*": Table S10

| **Functional category** | **Gene** | **Exon Coordinates** | **PSI**  **iPSC-CM** | **PSI Prenatal Hearts** | **PSI Postnatal Hearts** | **Affected Domains/ Motifs** |
| --- | --- | --- | --- | --- | --- | --- |
| Kinase | ***CLK1*** | chr2:200860125-200860215 | 0.38 | 0.95 | 0.97 | ORF disruption upon exon exclusion  Protein kinase domain (PF00069, PS50011) |
| Kinase | ***CLK4*** | chr5:178617344-178617434 | 0.17 | 0.96 | 1 | ORF disruption upon exon exclusion  Protein kinase dmain (PF00069, PS50011) |
| Cell Adhesion | ***ITGB1*** | chr10:32907063-32907143 | 0.22 | 1 | 1 | Integrin beta cytoplasmic domain (PF08725)  Phosphotyrosine binding domain  (ELME000122) |
| Methylation | ***METTL3*** | chr14:21499372-21499491 | 0.47 | 0.82 | 0.83 | ORF disruption upon intron inclusion  Disrupts MT-A70 domain (PF05063) |
| mRNA decay | ***PATL1*** | chr11:59655956-59656045 | 0.79 | 1 | 1 | Topoisomerase II-associated protein PAT1 (PF09770)  Proline-rich region (PS50099)  Highly disordered region |
| Cell cycle | ***PDE4DIP*** | chr1:149024445-149024684 | 0.13 | 0 | 0 | Highly disordered region |
| Splicing | ***RSRP1*** | chr1:25244454-25244498 | 0.54 | 0.02 | 0.02 | ORF disruption upon exon inclusion  Highly disordered region |
| Splicing | ***SNRPE*** | chr1:203862182-203862222 | 0.56 | 0.92 | 1 | LSM domain (PF01423) |
| Contraction | ***TPM1*** | chr15:63044027-63044152 | 0.72 | 0.93 | 0.95 | Tropomyosin domain (PF00261) |
| NMD | ***UPF3A*** | chr13:114286302-114286400 | 0.61 | 1 | 1 | Smg-4/UPF3 family (PF03467)  Glutamic acid-rich region (PS50313);  Disordered region |
