## Supplementary material for "Alternative splicing dynamics during human cardiac development *in vivo* and *in vitro*": Table S12

**Supplementary Table 10 –** Primers used for qRT-PCR validation of alternative splicing events.

| **Primer** | **Sequence** | **Type** | **Amplicon Size** |
| --- | --- | --- | --- |
| **Maturation-driven splicing events** | | | |
| *TNNT2* Ex5 Fw | GAGGACTGGAGAGAGGACGA | target AS event | 122 bp |
| *TNNT2* Ex7 Rv | GCCTCCTTTGCTTCCTCTTCT |  |  |
| *TNNT2* Ex14 Fw | AGCGGAAAAGTGGGAAGAGG | constitutive exon | 103 bp |
| *TNNT2* Ex14 Rv | AGCTGATCTTCATTCAGGTGGT |  |  |
| *CMYA5* Ex4 Fw | TCTGCAGAGCATGGACACTG | target AS event | 101 bp |
| *CMYA5* Ex4 Rv | TGATTTCCTCAAACGAAGTCAGG |  |  |
| *CMYA5* Ex8 Fw | ACAGTGAAAGAAAGCTACTGCA | constitutive exon | 107 bp |
| *CMYA5* Ex8 Rv | CTTTCACTGGGCAGGCTACA |  |  |
| *LDB3* Ex8 Fw | CTGCTTCAAGTCCTGCCGA | target AS event | 101 bp |
| *LDB3* Ex8 Rv | CTCACTGTAGCTGGTGTGGG |  |  |
| *LDB3* Ex6 Fw | AAGGACCTTGCCGTAGACAG | constitutive exon | 114 bp |
| *LDB3* Ex6 Rv | AGACTGCAGGTTGGAGGAAC |  |  |
| *TRDN* Ex19 Fw | GAACACTCAGTTCCAAGTGA | target AS event | 115 bp |
| *TRDN* Ex20 Rv | CTTTTTTAATTGAAACCGCA |  |  |
| *TRDN* Ex2 Fw | TGGATCTGTGCCCAAATCCC | constitutive exon | 95 bp |
| *TRDN* Ex2 Rv | TCAGGGCAATGACCAGAAGC |  |  |
| *CAMK2D* Ex17 Fw | TGCCAAAGACAATGCAGTCAG | target AS event | 201 bp |
| *CAMK2D* Ex18 Rv | AGACCCATATGTGAATGGTTTTCA |  |  |
| *CAMK2D* Ex10 Fw | GGACACGGTGACTCCTGAAG | constitutive exon | 102 bp |
| *CAMK2D* Ex10 Rv | ATCCATGGGTGCTTCAGTGC |  |  |
| **iPSC-CM splicing events** | | | |
| *CLK1* Ex4Fw | GGATGATGAGGAGGGTCACC | target AS event | 117 bp |
| *CLK1* Ex5Rv | TGATCGATGCACTCCACAACT |  |  |
| *CLK1* Ex9Fw | CGTGATGAACGCACCTTAATAA | constitutive exon | 116 bp |
| *CLK1* Ex9Rv | TGATCGATGCACTCCACAACT |  |  |
| *CLK4* Ex4Fw | GGATGATGAGGAGGGTCACC | target AS event | 110 bp |
| *CLK4* Ex5Rv | TGCACTCTACAACTTTGCCA |  |  |
| *CLK4* Ex2Fw | GCGGCATTCCAAAAGAACTCA | constitutive exon | 123 bp |
| *CLK4* Ex2Rv | TGCCTGTTCTCTTGTGTGCT |  |  |
| *SNRPE* Ex1Fw | GTGGCCAGGGTCAGAAAGTG | target AS event | 103 bp |
| *SNRPE* Ex2Rv | TCTGAAGATGAGGTTCTGCAC |  |  |
| *SNRPE* Ex5Fw | ACTCTGCTACAAAGTGTCTCCA | constitutive exon | 146 bp |
| *SNRPE* Ex5Rv | GCCATCTTGTAGTAACACGAGG |  |  |
| *METTL3* Int8Fw | GGGCCCAATTCAATAGGTGGA | target AS event | 143 bp |
| *METTL3* Ex9Rv | CTGGTTGAAGCCTTGGGGAT |  |  |
| *METTL3* Ex2Fw | CTACGGAATCCAGAGGCAGC | constitutive exon | 216 bp |
| *METTL3* Ex2Rv | CGTGGAGATGGCAAGACAGA |  |  |
